# Individualised computational modelling of immune mediated disease onset, flare and clearance in psoriasis

**DOI:** 10.1101/2021.09.18.460913

**Authors:** Fedor Shmarov, Graham R. Smith, Sophie C. Weatherhead, Nick J. Reynolds, Paolo Zuliani

## Abstract

Despite increased understanding about psoriasis pathophysiology, currently there is a lack of predictive computational models. We developed a personalisable ordinary differential equations model of human epidermis and psoriasis that incorporates immune cells and cytokine stimuli to regulate the transition between two stable steady states of clinically healthy (non-lesional) and disease (lesional psoriasis, plaque) skin. In line with experimental data, an immune stimulus initiated transition from healthy skin to psoriasis and apoptosis of immune and epidermal cells induced by UVB phototherapy returned the epidermis back to the healthy state. Notably, our model was able to distinguish disease flares. The flexibility of our model permitted the development of a patient-specific “UVB sensitivity” parameter that reflected subject-specific sensitivity to apoptosis and enabled simulation of individual patients’ clinical response trajectory. In a prospective clinical study of 94 patients, serial individual UVB doses and clinical response (Psoriasis Area Severity Index) values collected over the first three weeks of UVB therapy informed estimation of the “UVB sensitivity” parameter and the prediction of individual patient outcome at the end of phototherapy. An important advance of our model is its potential for direct clinical application through early assessment of response to UVB therapy, and for individualised optimisation of phototherapy regimes to improve clinical outcome. Additionally by incorporating the complex interaction of immune cells and epidermal keratinocytes, our model provides a basis to study and predict outcomes to biologic therapies in psoriasis.

**Author Summary:** We present a new computer model for psoriasis, an immune-mediated disabling skin disease which presents with red, raised scaly plaques that can appear over the whole body. Psoriasis affects millions of people in the UK alone and causes significant impairment to quality of life, and currently has no cure. Only a few treatments (including UVB phototherapy) can induce temporary remission. Despite our increased understanding about psoriasis, treatments are still given on a ‘trial and error’ basis and there are no reliable computer models that can a) elucidate the mechanisms behind psoriasis onset or flare and b) predict a patient’s response to a course of treatment (*e*.*g*., phototherapy) and the likelihood of inducing a period of remission. Our computer model addresses both these needs. First, it explicitly describes the interaction between the immune system and skin cells. Second, our model captures response to therapy at the individual patient level and enables personalised prediction of clinical outcomes. Notably, our model also supports prediction of amending individual UVB phototherapy regimes based on the patient’s initial response that include for example personalised delivery schedules (*i*.*e*., 3x weekly *vs*. 5x weekly phototherapy). Therefore, our work is a crucial step towards precision medicine for psoriasis treatment.

## 1 Introduction

Psoriasis vulgaris is a systemic immune-mediated inflammatory disease characterised by immune cell infiltration, keratinocyte hyperproliferation (up to an eight-fold increase in epidermal cell turnover) [8] and tortuosity of dermal blood vessels. Psoriasis is common (affecting 1-2% of “Western populations”) and manifests itself as red scaly plaques distributed over the whole skin, causing significant disability and impairment to quality of life [10]. As well as being associated with inflammatory arthritis in up to 30% of the patients, psoriasis is increasingly linked to metabolic syndrome and cardiovascular disease [42].

The pathophysiology of psoriasis is complex, multifactorial and thought to be triggered through environmental genetic interactions. For example, psoriasis can be triggered in response to innate or environment stimuli (*e*.*g*., injury or infection). It is known that both stimuli converge to myeloid dendritic cells and macrophages which then increase the production of cytokines such as IL-12 and IL-23 that stimulate the activation and proliferation of T-cell subsets, which in turn produce IL-17A, IL-17F, IL-20, IL-22, TNF and IFN-*γ* [59]. Tc17/Th17 represent the dominant T cell subset in lesional psoriasis, with expanded Tc17/Th17 clonotypes localising to psoriatic epidermis [48]. These cytokines stimulate keratinocyte growth and production of chemokines and keratinocyte-based growth factors further amplifying the immune response and maintaining the hyperplastic and chronic inflammation within psoriasis plaques [27, 35, 36, 37, 40, 15, 3]. The importance of structural cells in regulating the immune response in health and disease is increasingly recognised [34, 32]. Psoriasis is a chronic persistent disease but may undergo periods of remission; currently though there is no cure. Narrow-band ultraviolet B (UVB) phototherapy is one of just a few treatments that can clear psoriasis plaques over 8-12 weeks of therapy and induce a period of remission [33]; the treatment is usually prescribed to patients with mild to moderate psoriasis. As patients attend the hospital department 3 times weekly, this facilitates the regular recording of disease activity (Psoriasis Area and Severity Index [PASI] - a visual examination of psoriasis extent), clinical response (including disease exacerbations - flares) and any adverse events. Nevertheless, relapse is common (occurring in 70% of patients within 6 months from the end of the therapy); patients may then be considered for further UVB or switched to systemic therapy. At present, prediction of individual patient outcome to UVB or systemic therapy is not feasible, although some progress has been achieved [63]. For example, lower rates of psoriasis clearance in the early stages of the UVB therapy were negatively associated with the final clearance outcome [59].

Previous agent-based models [59] and ordinary differential equation (ODE)-based models [66] have simulated psoriasis onset and psoriasis clearance by induction of keratinocyte apoptosis through UVB phototherapy. However, despite the crucial role of acquired and innate immunity in initiating and maintaining psoriasis, these models provide only an implicit representation of the immune system.

The principal aims of the work presented in this paper were to develop a new ODE model of epidermis that: 1) explicitly describes the complex interplay between immune cells and keratinocytes; 2) maintains two stable steady states: a clinically healthy one (non-lesional skin) and a psoriatic one (lesional/plaque skin) and 3) enables switching between the two steady states via crossing of an unstable “transition” state through either the introduction of a sufficiently strong, transient immune stimulus (“healthy” to “psoriatic” transition) or by inducing an appropriate amount of apoptosis of proliferating keratinocytes and immune cells through UVB phototherapy (“psoriatic” to “healthy” transition). The inclusion of an immune component is important because it enables modelling the onset of psoriasis as well as flares. Furthermore, the immune component makes the model more generalisable, for example it would enable modelling the effects of biologics therapies, which are crucial for treating moderate to severe psoriasis. An additional important advance is the ability of our ODE model to take into account patient-specific “UVB sensitivity” through a corresponding parameter, and predict clinical outcome at the end of phototherapy based on the early clinical trajectory of response. (One of the factors contributing to a patient’s UVB sensitivity is measured clinically at the beginning of phototherapy (minimum erythema dose). Clearly, other factors (*e*.*g*., genetic ones) may influence a patient’s UVB sensitivity, but we do not explicitly model them in this work.) Together these results support the development of precision medicine in psoriasis.

## 2 Model and Methods

### 2.1 Epidermis model

Our model focuses on human epidermis (see Figures 1a and 1b) and incorporates the following cell species:

**Figure 1:**
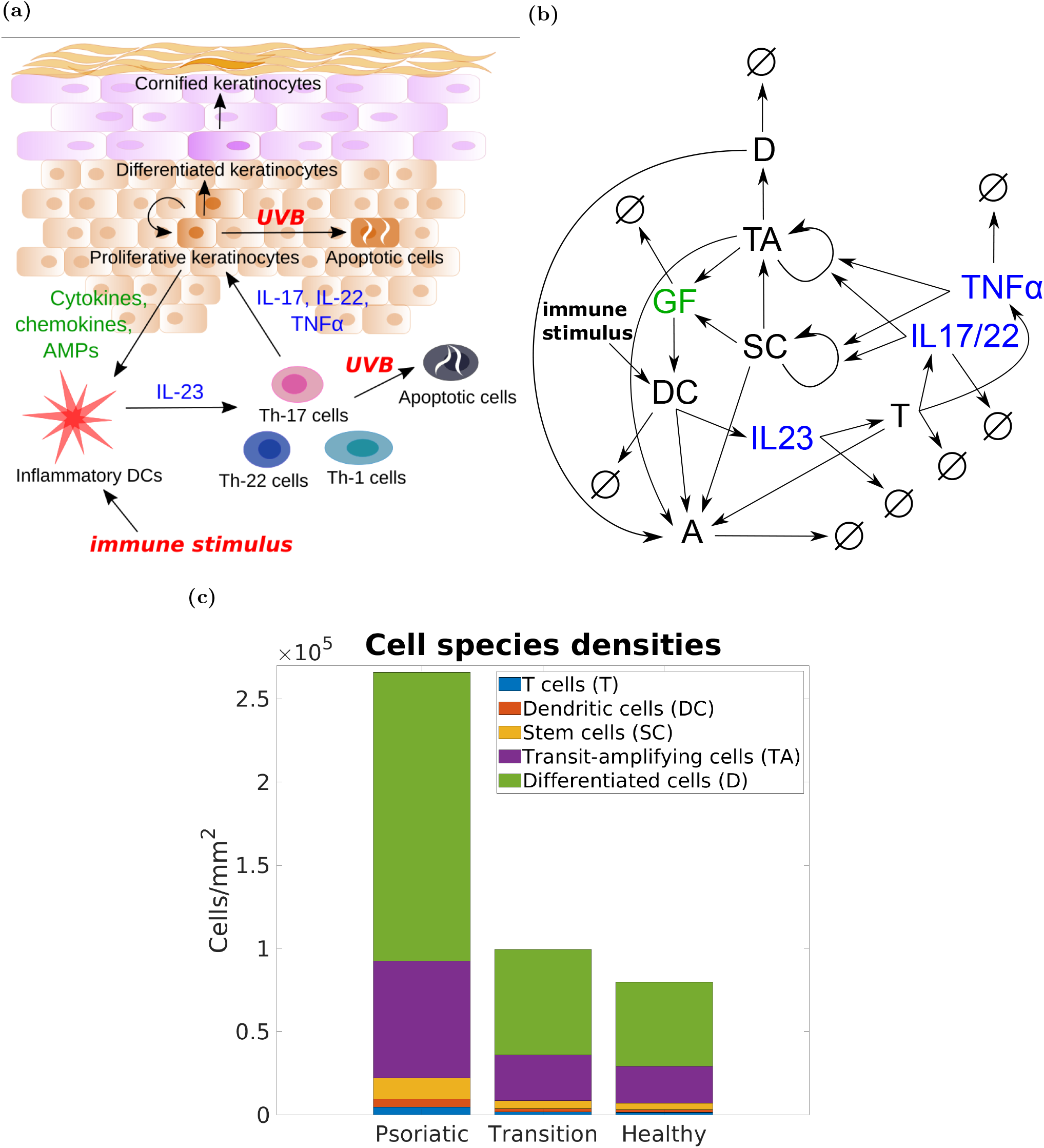
Our ODE model of epidermis describes an explicit interaction between the main types of keratinocytes and immune cells mediating psoriasis. Panels: (a) - mechanism of interaction between the main cell types and cytokines in epidermis; (b) - interactions between the species of our model, where *SC* - stem cells, *TA* - transit-amplifying cells, *D* - differentiating cells, *DC* - dendritic cells, *T* - T cells, *A* - apoptotic cells, *IL*_17*/*22_ - interleukin-17/22, *IL*_23_ - interleukin-23, *TNFα* - tumour necrosis factor alpha, *GF* - keratinocyte-derived growth factors, and ∅ - degradation species; (c) - cell density (cells*/mm*^2^) for every cell type (excluding apoptotic cells) in the healthy (non-lesional skin), psoriatic (lesional/plaque skin) and transition steady states.

- proliferating keratinocytes: stem cells (SC) and transit-amplifying cells (TA);
- differentiating keratinocytes (D).

The model also incorporates immune cells (T cells) and dendritic cells (DC), which may be located in either the epidermis or dermis but the location is not relevant to our model.

Proliferation and activation of SC and TA cells is mediated by IL-22, IL-17, type I interferons and TNF*α* cytokines produced by the activated T cells and/or dentritic cells [3]. In response to the increase in proliferation rate, keratinocytes increase their production of growth factors (GF) and chemokines that attract and activate dendritic cells producing more IL-23, which in turn activates more T cells, resulting in a positive feedback and self-sustaining loop (see Figure 1b).

In developing our model defined by the system of ordinary differential equations (ODEs) (1), the following three assumptions apply: 1) apoptosis induced by a single UVB dose lasts for 24 hours and affects proliferating keratinocytes and immune cells equally (see Section 2.2); 2) cell growth depends on cell density (see Section SM.5); and 3) we model clinically observable behaviour via a simplified PASI model that does not take the disease area into account (see Section 2.3). Our model is designed as a bi-modal switch consisting of three steady states: two are stable – clinically healthy (non-lesional skin) and psoriatic (lesional/plaque skin) states – and one is unstable (transition state). (The model features 32 steady states overall, but only the three states mentioned above lay in the positive real space: see Section SM.1 and Section SM.2.) The model design process considered the following main properties: epidermis composition, speed of psoriasis onset, epidermal turnover time, keratinocytes, apoptosis and desquamation rates. The specific parameters (apoptosis and desquamation rates, degradation of apoptotic cells, and production/degradation of cytokines and growth factors) were derived from published experimental and clinical data, and modelling outcomes described in the literature, as detailed below in Table 1. Notably, the ratio of production and degradation rate parameters for immune cells (T cells and DC) *vs*. cytokines in our model matches the ratio of half-life for CD4 T cells [56] and the IL-12 cytokine [7]. It is also important to note that the tissue levels of endogenous cytokines such as IL-23 are below the level of assay detection. Consequently, half-life measurements and pharmacokinetic models such as those reported in [13, 12] relate to exogenous administration of cytokines (or growth factors) and may therefore differ from our model which includes a mass-action dependence of IL-23 production on dendritic cells (or dependence on stem cells and TA cells for growth factor production) – we thus assume comparable values for the cytokines and growth factors parameters in our model.

**Table 1:** Model parameters.

| Name | Description | Value | Units | Source |
| --- | --- | --- | --- | --- |
| $il_{1720}$ | IL-17 degradation rate constant | 36.5 | $d^{-1}$ | inferred from [7, 56] |
| $il_{2320}$ | IL-23 degradation rate constant | 36.5 | $d^{-1}$ | inferred from [7, 56] |
| $tnf_{20}$ | TNF degradation rate constant | 36.5 | $d^{-1}$ | inferred from [7, 56] |
| $gf_{20}$ | GF degradation rate constant | 36.5 | $d^{-1}$ | inferred from [12] |
| $il_{232}$ | IL-23 production rate constant by dendritic cells | 1 | $a.u./(cells \times d)$ | inferred from [13] |
| $il_{172}$ | IL-17 production rate constant by T cells | 0.5 | $a.u./(cells \times d)$ | inferred from [13] |
| $tnf_2$ | TNF production rate constant by T cells | 0.5 | $a.u./(cells \times d)$ | inferred from [13] |
| $sc_{2gf}$ | GF production rate constant by SC | 1 | $a.u./(cells \times d)$ | inferred from [12] |
| $ta_{2gf}$ | GF production rate constant by TA | 1 | $a.u./(cells \times d)$ | inferred from [12] |
| $t_2$ | T cells activation rate constant | 55 | $cells/(a.u. \times d)$ | inferred from [51] |
| $t_{20}$ | T cells deactivation rate constant | 1.51 | $d^{-1}$ | inferred from [51, 9, 56] |
| $dc_{20}$ | Deactivation rate constant of DC | 1.51 | $d^{-1}$ | inferred from [9, 56] |
| $p$ | Coefficient for limiting keratinocytes growth | 2 | - | assumed |
| $dc_{vm}$ | Maximum activation rate constant of DC | 6000 | - | assumed |
| $dc_{kp}$ | Dissociation constant in Hill equation for DC | $3 \cdot 10^{12}$ | - | assumed |
| $dc_{act}$ | Basal activation rate constant of DC | 1880 | $cells/d$ | inferred from [9] |
| $n$ | Hill coefficient for DC | 4 | - | assumed |
| $sc_2$ | SC production rate constant | 0.0017635 | $(d \times a.u.)^{-1}$ | inferred from [60, 51] |
| $sc_{2ta}$ | TA production rate constant by SC | 0.0015415 | $(d \times a.u.)^{-1}$ | inferred from [44] |
| $sc_{20}$ | Proliferation limiting factor for SC | 1.8135e-05 | $d^{-p}$ | inferred from [44, 59] |
| $ta_2$ | TA production rate constant | 0.0017562 | $(d \times a.u.)^{-1}$ | inferred from [45] |
| $ta_{20}$ | Proliferation limiting factor for TA | 3.7321e-06 | $d^{-p}$ | inferred from [45] |
| $ta_{2d}$ | Differentiation rate constant | 0.0018063 | $(d \times a.u.)^{-1}$ | inferred from [60] |
| $d_{20}$ | Differentiation limiting factor | 5.05e-07 | $d^{-p}$ | inferred from [60] |
| $d_{desq}$ | Desquamation rate constant | 0.0047529 | $d^{-1}$ | [39] |
| $a_{base}$ | Basal apoptosis rate | 0.0036 | $d^{-1}$ | [59, Figure 1b] |
| $a_{20}$ | Degradation rate of apoptotic cells | 36 | $d^{-1}$ | [58, Figure 5-5] |

Each parameter was tested in turn to ensure the model remained stable, and some of the assumed parameters were adjusted in line with published data to ensure the model bistability. Also, due to lack of clinical and biomarker data, the values of the parameters of Table 1 have not been subject to personalisation. Should data be available, it would be possible to personalise further the model in terms of the assumed parameters.

In the next three subsections we describe in detail how our epidermis model is valid with respect to general clinical and biological measurements. In Section 3.3 we describe validation of our PASI and UVB phototherapy modelling with respect to individual PASI trajectories of our patient cohort.

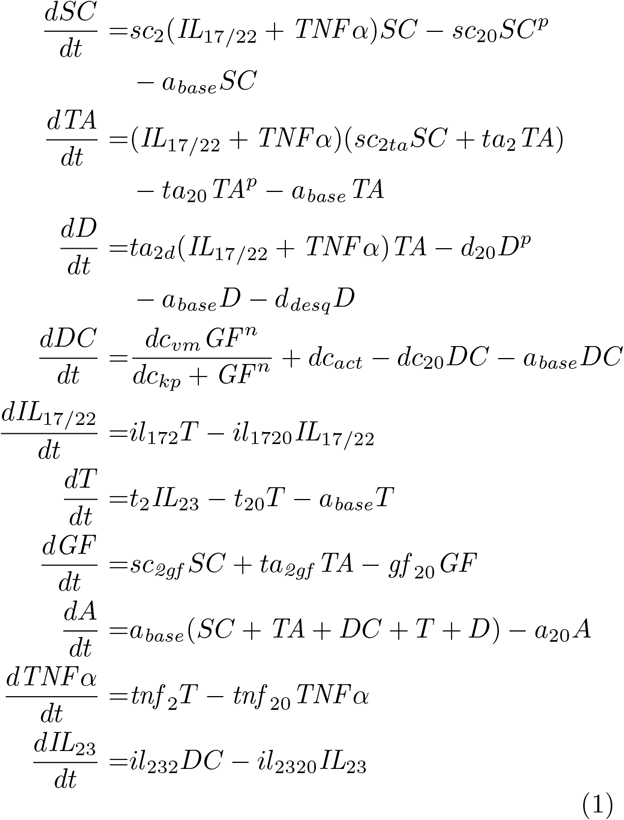

#### 2.1.1 Epidermis composition

Figure 1c shows the epidermis composition in the three steady states of our model (the precise values are reported in Table 2). The total number of cells per unit area of human epidermis is different from person to person, and varies across the human body. Ref. [9] reports a total cell density of 73,952 ± 19,426 *cells/mm*^2^ (mean and standard deviation) and 1,394 ± 321 *cells/mm*^2^ for Langerhans cells (*i*.*e*., dendritic cells in epidermis). In line with these data, the total cell and Langerhans cells densities in our model are 79,828 *cells/mm*^2^ and 1,563 *cells/mm*^2^, respectively.

**Table 2:** Steady states of the model

| Species | Description | Healthy<br>(non-lesional) | Transition | Psoriatic<br>(lesional/plaque) | Units |
| --- | --- | --- | --- | --- | --- |
| $SC$ | Stem cells | 3,947.21 | 4,855.46 | 12,544.23 | $cells/mm^2$ |
| $TA$ | Transit-amplifying cells | 22,224.37 | 27,308.71 | 70,349.64 | $cells/mm^2$ |
| $D$ | Differentiating cells | 50,529.36 | 63,458.55 | 173,504.14 | $cells/mm^2$ |
| $T$ | T cells | 1,556.09 | 1,897 | 4,782.97 | $cells/mm^2$ |
| $DC$ | Dendritic cells | 1,563.06 | 1,905.5 | 4,804.4 | $cells/mm^2$ |
| $A$ | Apoptotic cells | 7.98 | 9.94 | 26.6 | $cells/mm^2$ |
| $\Sigma$ | Total cell count | 79,828.07 | 99,435.16 | 266,011.98 | $cells/mm^2$ |
| $IL_{17/22}$ | Interleukin 17/22 | 21.32 | 25.99 | 65.52 | $a.u.$ |
| $IL_{23}$ | Interleukin 23 | 42.82 | 52.20 | 131.63 | $a.u.$ |
| $TNF$ | Tumor necrosis factor alpha | 21.32 | 25.99 | 65.52 | $a.u.$ |
| $GF$ | Keratinocyte-based growth factors | 717.03 | 881.21 | 2,271.06 | $a.u.$ |

The proportions of cell species in the healthy steady state are consistent with the values found in the literature. From Table 2 we see that our model features 63.2% of differentiating cells (40-66% is reported by [59]), 27.8% of transit-amplifying cells, 4.9% of stem cells, 1.8% dendritic cells and 1.8% of T cells. Hence, our model suggests that up to 96% of all cells in the epidermis are keratinocytes [59, 51].

#### 2.1.2 Psoriasis onset

We model the onset of psoriasis by increasing the activation rate of dendritic cells (corresponding to parameter *dc*_*act*_ in the equation for model species *DC* in (1)) by a constant amount *dc*_*stim*_ for *τ*_*stim*_ days. This produces psoriasis onset at different rates: from fast (as quick as 10 days) to slow (as long as 14 weeks). That is consistent with the known time lags between the onset of an immunological stimulus (*e*.*g*., streptococcal sore throat) and the first appearance of psoriatic lesions [59, 54, 28].

In the psoriatic state the growth rate of the proliferating keratinocytes (SC and TA cells) and their differentiation rate increase with respect to the healthy state by 9.74 and 9.73 times, respectively (see Section SM.4 for more details). Compared to normal skin, the increased number of proliferating keratinocytes *in vivo* ranges in psoriasis from 2-3 times [14, 48] to 6-8 times [8, 19]. As a result, the cell density in the psoriatic state of our model is 266,012 *cells/mm*^2^, *i*.*e*., 3.33 times the density in the healthy state, which is within the 2-5 times range reported in clinical studies [55, 61].

#### 2.1.3 Epidermal turnover time

In psoriasis, despite an increase in epidermis thickness, the epidermal turnover time drops by up to 4-7 times [62, 30]. This implies that the cell growth rate cannot linearly depend on the cell density (see Section SM.5). Thus, in each ODE, we introduced the non-linear term *a* · *X*^*p*^, where *a* is a constant, *X* is the corresponding keratinocyte species, and *p* is a parameter modelling the nonlinear dependency between the keratinocyte growth rate and the cell density. Parameter *p* is the “growth limiting constant” and models factors that influence the keratinocytes growth. (The higher the value of *p* the higher the ratio between the turnover times of the psoriatic and healthy states - for more details see SM.5.) For simplicity, we assumed *p* = 2, but any *p* > 1 can be used for personalising the model.

For setting the parameter values related to the growth rate, we applied epidermal turnover time values reported in the literature. Turnover times for healthy epidermis vary from 39 days [60], to 47-48 days [31] to 40-56 days [30], while a previously published model [66] predicts 52.5 days.

Similarly to [60, 66], the epidermal turnover time *τ* in our model can be calculated as the sum of the turnover time in the proliferating and the differentiating compartments using the formula:

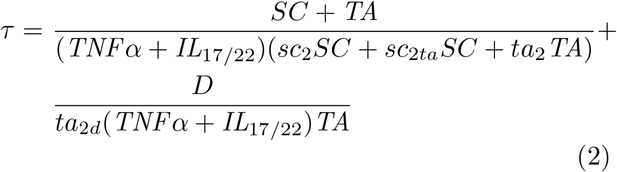

where *TNFα* is the tumour necrosis factor *α*, and the cell species densities are taken from either the psoriatic state or the healthy state (see Table 2). In our model, the epidermal turnover time in psoriatic epidermis drops almost three-fold in comparison to healthy epidermis, from 41.31 days to 14.25 days, consistent with *in vivo* studies [62, 30].

#### 2.1.4 Apoptosis and desquamation rates

Cell apoptosis and desquamation (terminal differentiation) are the two main mechanisms in our model by which keratinocytes can leave the proliferating and the differentiating compartments. It is technically challenging to measure rates and duration of apoptosis in human tissue, and so the parameters used in the model are based on data from mouse skin and *in vitro* studies of human keratinocytes [58, 29, 38, 59, 39]. In addition, it is assumed that in healthy epidermis all cells undergo apoptosis at the same rate (relative to their cell mass). In Section SM.6 we provide more details.

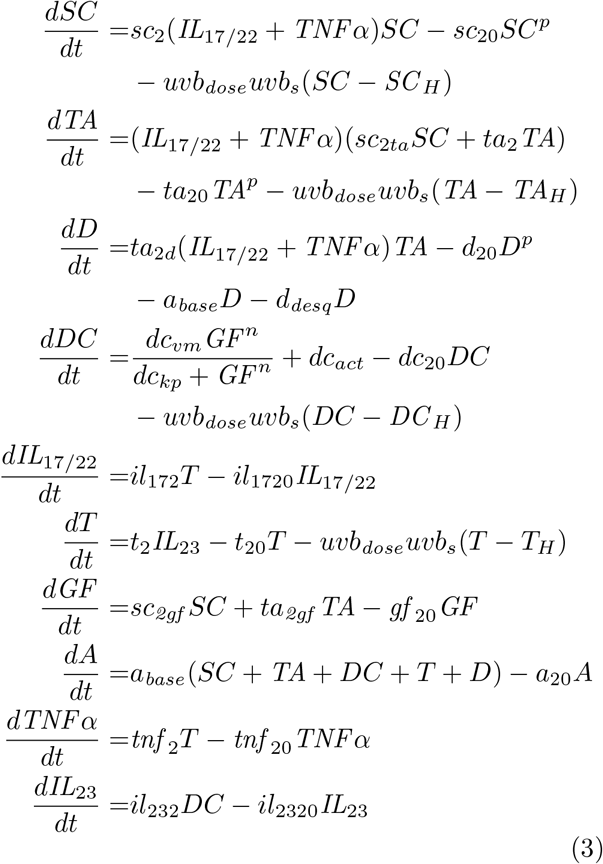

### 2.2 Modelling UVB phototherapy

In our model, psoriasis clearance with UVB is implemented by increasing the rate of apoptosis of proliferating keratinocytes (*SC* and *TA*) and immune cells (*DC* and *T*) for *t*_*uvb*_ hours by *uvb*_*dose*_·*uvb*_*s*_·(*X*−*X*_*H*_), where *uvb*_*dose*_ is the administered UVB dose (in *J/cm*^2^), *X* is the target species cell density, *X*_*H*_ is the target species cell density in the healthy steady state, and *uvb*_*s*_ is the “UVB sensitivity” parameter in (*J*^−1^*d*^−1^). Note that differentiated keratinocytes (*D*) may undergo apoptosis when UVB irradiated, but at a negligible rate [58, Section 5.3.4]. Hence, our model does not increase the apoptotic rate of differentiated cells during UVB irradiation. In Eq. (3) we give the full ODEs for our model when simulating UVB phototherapy as explained above. (The constants *SC* _*H*_, *TA*_*H*_, *DC* _*H*_ and *T*_*H*_ are found in Table 2, column “Healthy”.)

We assume that UVB induced apoptosis is distributed between 0 and 24 hours, as the peak of cell apoptosis is reported to be between 18 and 24 hours following UVB irradiation [59, Figure S2], *i*.*e*., we assume *t*_*uvb*_ = 24 hours. (If needed, the model can be reparameterised to allow for a shorter or longer duration of apoptosis.) Thus, the rate of UVB induced apoptosis is proportional to the applied dose (defined in terms of energy per unit area), current cell density (*i*.*e*., the higher the cell density the higher the number of apoptotic cells) and the patient-specific parameter UVB sensitivity (larger sensitivity values result into higher apoptosis rates). The term (*X* − *X*_*H*_) is introduced to account for the fact that the thickness of healthy epidermis does not reduce in response to UV [47, 46]. We also restrict our model to operate only between the healthy and psoriatic states thus, the term (*X* − *X*_*H*_) is always kept positive. Standard clinical protocols administer narrow-band UVB three times a week (*e*.*g*., Monday, Wednesday and Friday) for up to 12 weeks, with increasing graduated dose increments (on alternate doses) over the course of the treatment. An example of UVB dosimetry is shown in Figure 2a. The changes in cell densities and cytokine concentrations following the indicated UVB regime are shown in Figure 2b and Figure 2c; the rate of cell apoptosis is shown in Figure 2d. The rate of cell apoptosis in the model falls back to the basal value shortly after the end of the 24-hour period. This is because apoptotic cells are removed from the epidermis fairly quickly in our model (mean lifetime of 40 minutes – see Section SM.6). The obtained apoptosis values are consistent with those reported *in vivo* [59, Figure 1e].

**Figure 2:**
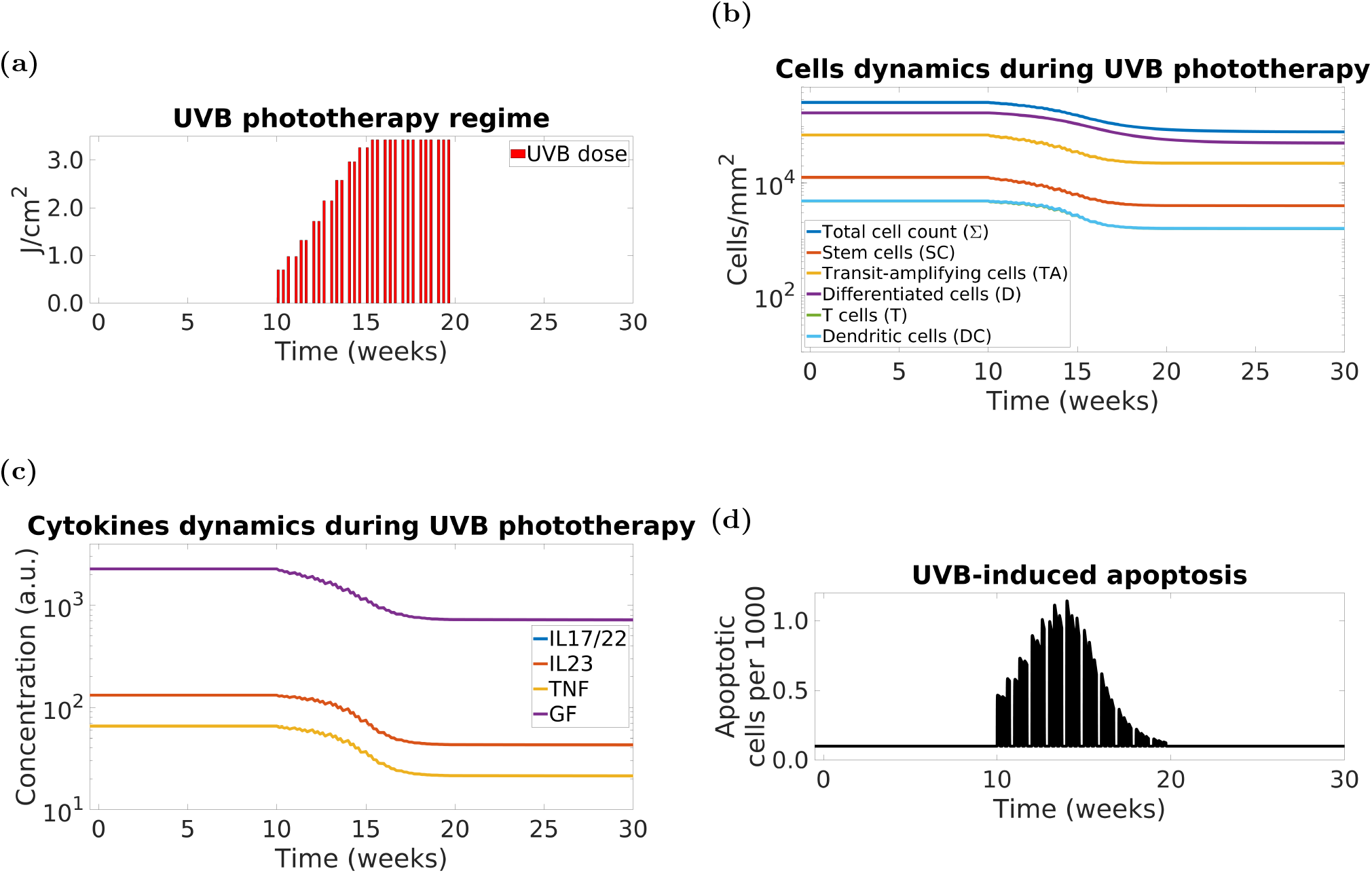
ODE psoriasis model demonstrates that UVB-induced apoptosis leads to psoriasis clearance. Simulation of the model response to a 10-week UVB phototherapy. Panels: (a) - simulated UVB irradiation regime, (b) - cell densities, where *SC* - stem cells, *TA* - transit-amplifying cells, *D* - differentiating cells, *DC* - dendritic cells, *T* - T cells, and Σ = *SC* + *TA* + *D* + *DC* + *T* - total number of cells cell density, (c) - cytokines concentration, where *IL17 /22* - interleukin-17/22, *IL23* - interleukin-23, *TNF* - tumour necrosis factor alpha, and *GF* - keratinocyte-derived growth factors, (d) - number of apoptotic cells per 1,000 cells.

Our model predicts complete clearance and eventual remission when the model dynamics drops below the “transition” steady state (*i*.*e*., at approximately 90% clearance). These data are consistent with recent findings in a prospective study of 100 patients in which achievement of PASI90 (*i*.*e*., at least 90% improvement over their initial PASI) pointed to longer remission [57]. However, not all patients achieving PASI90 progress to complete clearance and/or remission.

#### Apoptosis *vs*. Growth Arrest

Psoriatic epidermis was thought to be resistant to apoptosis [64] although only few previous studies have investigated before and during the early stages of therapy. Our experimental work [59, 1] provides clear evidence that therapeutic UVB irradiation induces apoptosis of psoriatic epidermis within 24 hours of irradiation. Our previous agent-based modelling suggested that apoptosis was sufficient to account for epidermal remodelling during resolution of psoriasis. UVB also induces growth arrest of cultured keratinocytes [6] and epidermal keratinocytes in normal human skin [43] but whether UV-induced growth arrest of immune cells or keratinocytes contributes to psoriasis plaque clearance remains unknown. We thus considered whether in our model growth arrest could account for clearance during UVB therapy. We found that growth arrest alone (no apoptosis) could induce clearance with a delay of several weeks after the end of the therapy (see Figure S4 – similar results are reported by [59]). However, in clinical practice psoriasis does not improve after UVB treatment completion, and therefore we do not further consider growth arrest in this paper.

### 2.3 Modelling PASI

The Psoriasis Area and Severity Index (PASI) is used for assessing the severity of the ongoing disease [65, 4, 50]. It ranges from 0 to 72, and it is calculated as a weighted sum of sub-scores corresponding to four body regions:

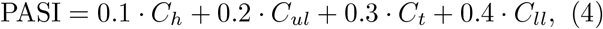

where *C*_*h*_, *C*_*ul*_, *C*_*t*_ and *C*_*ll*_ are the sub-score values for head, upper limbs, trunk and lower limbs, respectively. Each sub-score is obtained as

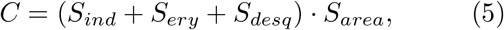

where *S*_*ind*_, *S*_*ery*_ and *S*_*desq*_ are values between 0 and 4 representing the severity of induration (thickness), erythema (redness) and desquamation (scaling), respectively, for the four body regions; *S*_*area*_ is a value between 0 and 6 representing the extent of the affected area.

We mimic patients’ PASI by using the species of our ODE model as follows: we use the total cell density

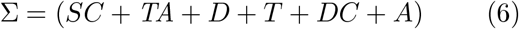

as a proxy for epidermal thickness. As we scale between 0 and 1 all the species modelling the PASI components (see below), we use the T cell density to represent inflammation which translates to erythema clinically within the PASI score for simplicity. The scaled dynamics of the cytokine species (*i*.*e*., IL-17, IL-23 and TNF*α*) is virtually identical to the scaled dynamics of T cells as these two families of species share the same shape of equation – a standard mass-action law (see the ODE model (1)) – and both results in scaled densities very close to 0, hence we only use T cell dynamics to estimate erythema. To assess scaling we use the non-proliferating cells *D* density.

In our model we do not take the surface area into account, and we only consider severity of PASI components over a unit area (*mm*^2^). Thus, our model assumes that the affected area stays constant throughout the therapy, while all the other components change (*i*.*e*., the plaques “fade away” rather than “shrink” in size).

If we rescale the species concentration to the [0, 1] interval (where 0 and 1 represent healthy and psoriatic steady states, respectively), we can model the relative change in each of the psoriasis symptoms *S*_*ind*_, *S*_*ery*_ and *S*_*desq*_ over a unit area of skin. After weighting the rescaled species we calculate the severity of the symptoms over a unit area for each body region (*i*.*e*., an estimate for *S*_*ind*_ + *S*_*ery*_ + *S*_*desq*_) as

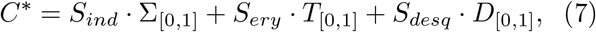

where Σ_[0,1]_, *T*_[0,1]_ and *D*_[0,1]_ are the cell species rescaled to [0,1]. Thus, obtaining *C*^*^ for each body region, and assuming that the amount of energy delivered by UVB per unit of area is the same across the entire surface of the body, the resulting PASI value can be modelled by

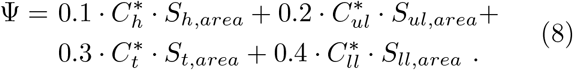

However, PASI subscores are seldom recorded in clinical practice – only the cumulative score is likely to be available. In order to mitigate this issue, we consider the average behaviour of the components over the entire body instead of modelling each body region separately. As a result, we introduce a new species to simulate a patient’s PASI when the PASI subscores are not available:

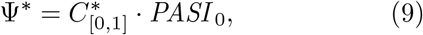

where 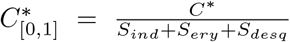 is the value of *C*^*^ rescaled to [0,1], and *PASI* _0_ is the baseline PASI. In the data we used in this work the PASI subscores are not available, and we thus used Ψ^*^ instead of Ψ and we assumed *S*_*ind*_ = *S*_*ery*_ = *S*_*desq*_ = 1 in all our experiments (in other words, we assumed the severity of induration, erythema, desquamation are all equal (to 1) in our model).

### 2.4 Model personalisation

#### Patient data

Our clinical data are derived from a prospective cohort of 94 patients receiving narrow-band UVB therapy for psoriasis, recruited at the Royal Victoria Infirmary, Newcastle upon Tyne (UK). The dataset is described fully in [57] but in brief it includes serial weekly patients’ PASI measurements (median 7; range 4-11), corresponding serial UVB doses (median 24; range 4-11), delivered 3 times a week together with data on time to relapse and PASI at relapse, collected for up to 18 months. Out of 94 patients, 6 subjects did not relapse within the 18-month monitoring period, and 26 individuals were lost to follow-up.

#### Parameter fitting

We estimate the UVB sensitivity parameter *uvb*_*s*_ to fit the model PASI simulations to the patients’ PASI data. For a given patient, the value of their *uvb*_*s*_ parameter is the *minimal uvb*_*s*_ value that minimize

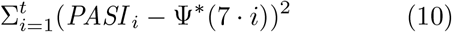

where *t* is the number of points (weeks) in the patient’s PASI trajectory (the model time units are days while patients data are recorded weekly, hence the use of 7 · *i* in Ψ^*^) and *PASI* _*i*_ is the (single) PASI measurement at the end of week *i* of phototherapy. The minimisation of (10) (as a function of *uvb*_*s*_) while also minimising *usb*_*s*_ is performed via an exhaustive search over the [0, 1] interval, starting at 0 with a step of 0.01. For every value of *uvb*_*s*_ we simulate our ODE model (1) with Ψ^*^ defined by (9) and Ψ^*^(0) = *PASI* _0_ and all other model species set at the psoriatic steady state. We then compute the error given by (10) and update the ‘minimum error’ and the ‘minimum’ *uvb*_*s*_ variables if the newly computed error is lower than current minimum. This method clearly allows minimising the value of *uvb*_*s*_ since from all the parameter values producing the lowest value for the objective function (10) we choose the lowest value for *uvb*_*s*_. The minimisation of *uvb*_*s*_ is important because it prevents the model from being ultra responsive to treatments via unrealistically large *uvb*_*s*_ values, which would represent patients that are overly sensitive to small doses of UVB, such as a very small fraction of their MED perhaps comparable to an average daily exposure to sunlight. Paradoxically, UVB does not induce significant erythema in lesional (plaque) skin. Localised irradiation with multiple (x8 or x16) MEDs can be directed to localised plaques through 308 nm lasers for example [5] resulting in rapid clearing. However, as such doses given as whole body irradiation would cause significant erythema and burning of non-lesional skin, the parameters within the model constrain psoriasis clearance from occurring with a small number of high MED doses of UVB irradiation. In the clinical dataset used for this work, the smallest recorded number of doses to achieve complete clearance (PASI100) was 18, *i*.*e*., six weeks of UVB phototherapy.

#### Flares

During longitudinal follow up [23, 18] and during the course of the therapy some patients may experience spontaneous disease exacerbations or flares (*i*.*e*., worsening of the symptoms and signs due to undefined environmental/immunological stimuli). We simulate patients’ flares by introducing and fitting parameters *dc*_*stim,i*_ for *i* = 1, …, *t* (where *t* is the number of PASI values in the patient’s trajectory) for every patient. Each such parameter models an immune stimulus of constant strength lasting 7 days, since in our dataset we have at most one PASI reading per week. (These parameters will increase the activation rate of dendritic cells from their basal value *dc*_*act*_ – see the equation for species *DC* in Eq. (1).) The parameters *dc*_*stim,i*_ are fitted sequentially beginning with *i* = 1, and every parameter is searched incrementally in the interval [0, 6,000] starting at 0 with a step size of 60 (*i*.*e*., 100 increments). The current parameter value is increased until the objective function value (10) starts rising. This parameter value is set in the model, and the fitting of the next parameter *dc*_*stim,i*+1_ is performed.

#### Statistical methods

The goodness of fit of our model is assessed by calculating the distribution of the difference (*PASI* _*i*_ − Ψ^*^(7 · *i*)) between the patients’ PASI trajectories and the model PASI simulation for all but the baseline PASI values. (Our dataset contains 754 PASI values distributed between 0 and 24.1, with mean 3.41, median 2.3 and IQR [1, 4.5]). We obtain the distribution of the PASI estimation error by uniformly sampling (*PASI* _*i*_ − 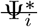) for each patient. We repeat this process 1,000 times and compute mean, median and standard deviation of the resulting cumulative distributions, which are then compared with PASI assessment errors made by formally trained physicians [22].

## 3 Results

We developed an ODE model of normal and psoriasis skin that would enable direct comparison to patient specific disease trajectories and prediction of outcomes to therapy at an individual patient level. As outlined in Section 2.1, our model’s behaviour (including key indicators such as proliferation rates and epidermal turnover time), is consistent with data from clinical studies. In the sections reported below, we systematically studied the dynamic behaviour of our model for predicting the speed of psoriasis onset and the impact of varying UVB therapy regimes and UVB sensitivity to outcome. Finally, we explored the capabilities of our model for personalisation of disease trajectories and for stratification of patients undergoing disease flares.

### 3.1 ODE model suggests speed of psoriasis onset depends on strength and duration of immune stimulus

As described in Section 2.1, the onset of psoriasis is initiated by an immune stimulus that increases the activation rate of the dendritic cells. Our analysis indicates that the speed of psoriasis onset depends on both the strength and the duration of this immune stimulus (Figure 3). When the immune stimulus is sufficient to drive the system through the transition state, the model will inevitably progress to the psoriasis steady stats. In order to provide confidence that this transition has resulted in psoriasis, to generate Figure 3 we have set the threshold relatively high along the transition path at 90% of the distance between normal and fully developed psoriasis state.

**Figure 3:**
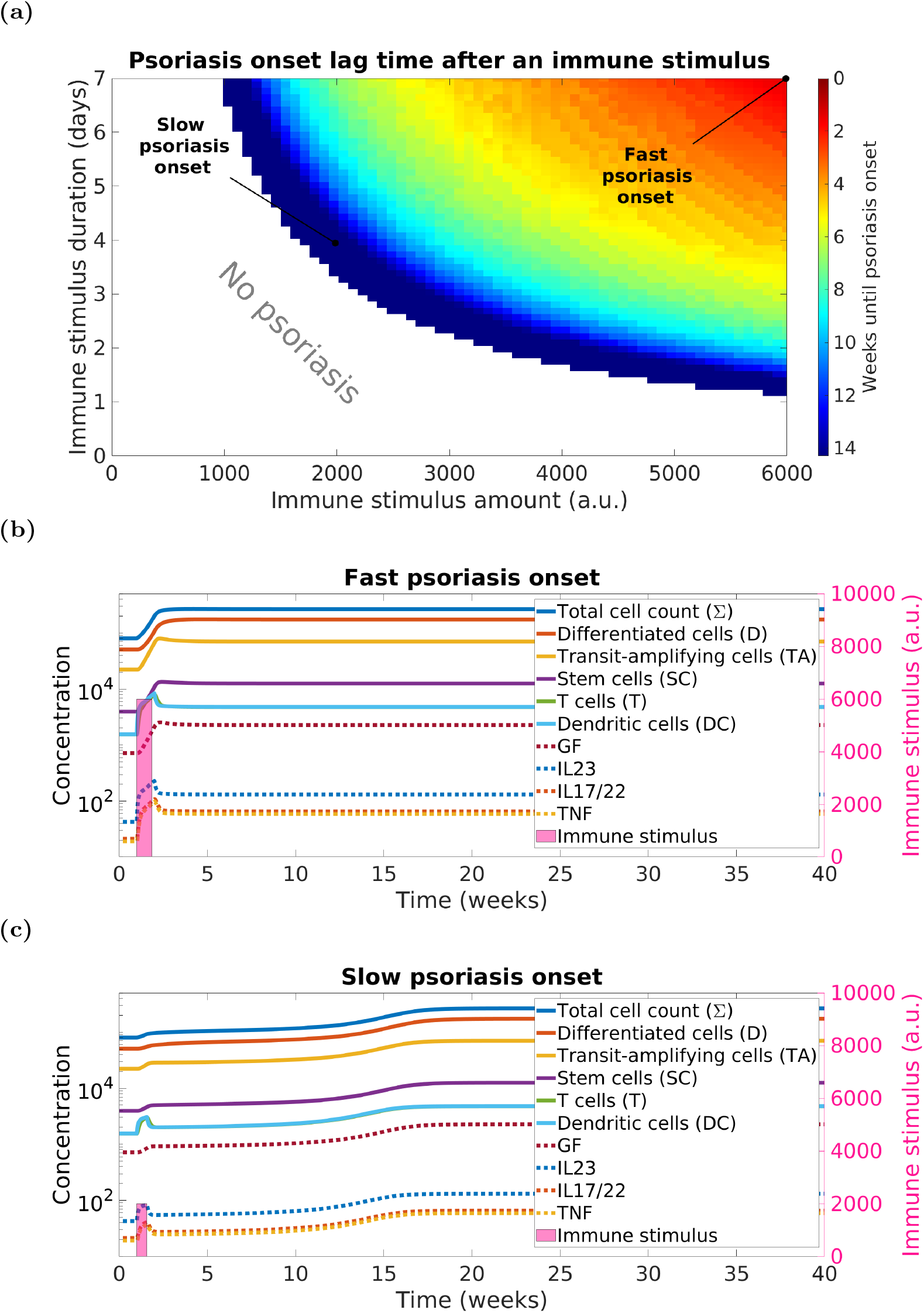
The speed of psoriasis onset in ODE model depends on the strength and the duration of the immune stimulus. Panels: (a) - heatmap where white-coloured area denotes combinations of immune stimulus strength and duration that do not lead to psoriasis; other colours denote psoriasis; (b) and (c) - examples of model simulations for the combinations of stimulus strength and duration values, as highlighted in Panel (a), leading to fast and slow psoriasis onsets, respectively. Psoriasis onset occurs when *totC* = *totC*_*H*_ + 0.9(*totC* _*P*_ − *totC* _*H*_) ≈ 247,376 *cells/mm*^2^ ≈ 0.93 · *totC*_*P*_ (*i*.*e*., the total cell density of the model has covered 90% of the distance between the healthy state and the psoriatic state – see Table 2 for the actual cell densities). This is due to the relatively slow convergence of the model to the psoriatic steady state.

In addition, Figure 3 demonstrates the model dynamics for two exemplary scenarios simulating slow (14 weeks) and fast (10 days) psoriasis onset. Within the ranges explored for stimulus strength and duration, 14 weeks is the longest delay and 10 days the shortest delay to a *full* psoriasis plaque achievable by our model – clinical observations report psoriasis onset no sooner than 4 days after an immune stimulus [28].

### 3.2 ODE model simulates individual clinical outcomes and personalised amendments of phototherapy to induce psoriasis clearance and remission

Our model indicates that a minimum number of UVB episodes and appropriate irradiation frequency are necessary for clearing psoriasis and inducing remission, depending on the patient-specific UVB sensitivity parameter *uvb*_*s*_ (*i*.*e*., patient-specific UVB sensitivity to phototherapy), and actual UVB doses that will be administered.

Following a given UVB irradiation regime, our model can simulate different outcomes in which the chances of psoriasis clearance increase for higher values of *uvb*_*s*_. For example, two models with different UVB sensitivity values (modelling two patients) receiving equal amounts of UVB might not reach the same outcome. The heatmaps presented in Figure 4 depict the model outcome as a function of the number of UVB doses (of a given therapy regime) and the UVB sensitivity parameter *uvb*_*s*_. Figures 4a and 4b show that changing therapy from 3 times to 5 times a week can, overall, increase the likelihood and speed of psoriasis clearance, consistent with results from a (small) clinical trial [17] and a recent survey [16]. However, as in [17], the actual improvement is modest and might not be clinically justifiable because of inconvenience or risk of side effects (*e*.*g*., erythema or “sun burn”). A standard protocol for UVB phototherapy is treatment three times per week with a minimum of 24 hours between sessions but clinical studies and our modelling indicates (see Section SM.7) that lower frequencies of irradiation (*e*.*g*., twice weekly [11]) may also be effective although may take longer in absolute time to achieve clearance.

**Figure 4:**
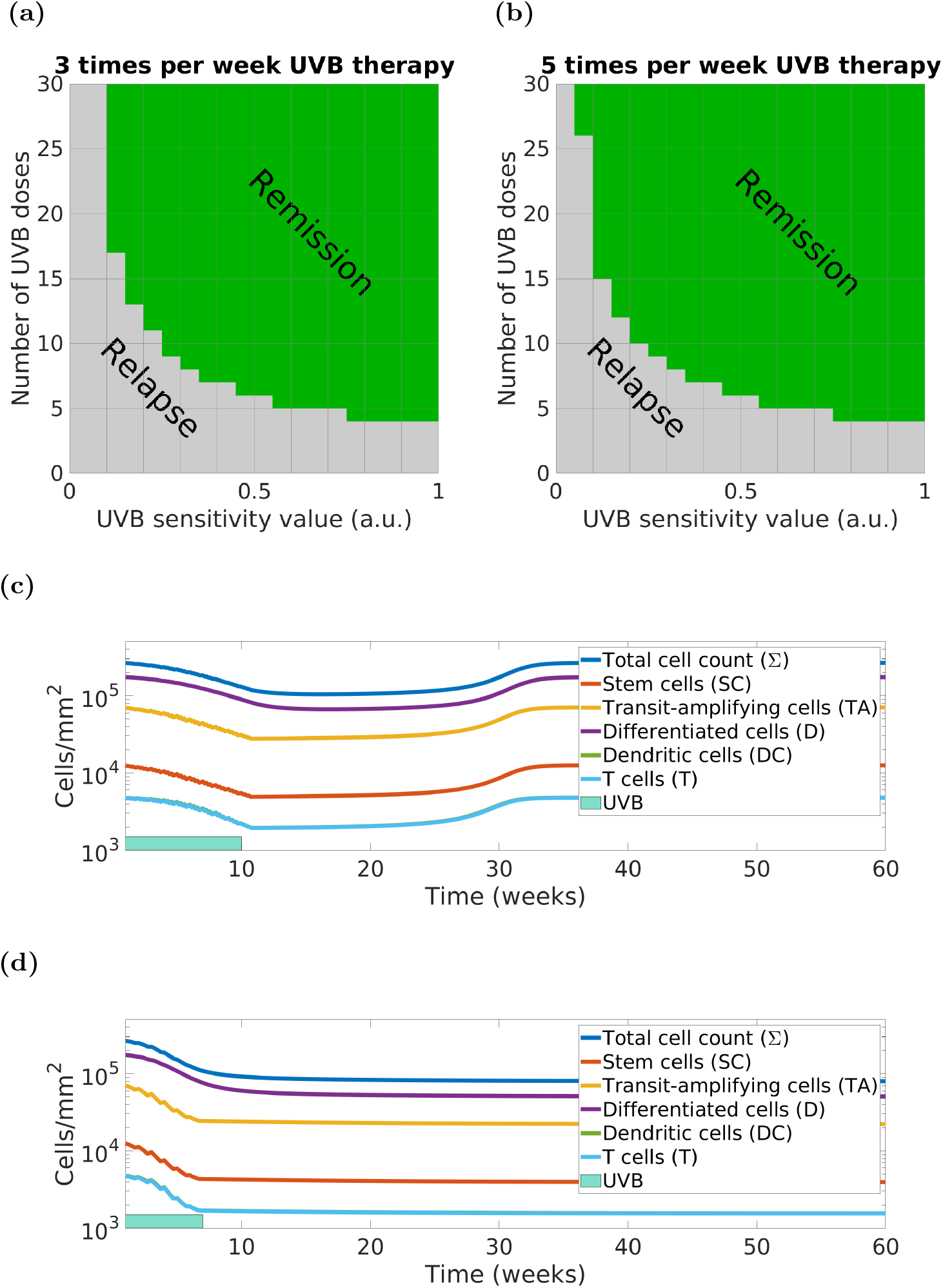
Our ODE model predicts that the total number of UVB doses, irradiation frequency and the patient’s UVB sensitivity (modelled by the *uvb*_*s*_ parameter) are the key factors in designing a personalised therapy for achieving psoriasis clearance and remission. Panels: (a) and (b) - green areas contain the values for which the model predicts remission following the therapy; (c) and (d): personalised model simulation of a virtual patient (*uvb*_*s*_ = 0.05) whose 3x weekly phototherapy (c) fails to clear psoriasis while 5x weekly phototherapy (d) with the same doses (*n* = 30) succeeds.

If within Figure 4, we consider a patient characterised by a “low” *uvb*_*s*_ = 0.05, we can then compare the model simulations of a therapeutic phototherapy course consisting of 30 doses but delivered 3 times *vs*. 5 times a week and how this affects relapse. The 3 times a week simulation (Figure 4c) results in relapse of psoriasis within a few months, whereas the 5 times a week therapy regime (Figure 4d) induces a longer duration of remission, thereby showing that some patients might potentially benefit from phototherapy delivery adjustments.

Finally, we note that the *uvb*_*s*_ parameter is associated with the rate of keratinocyte and lymphocyte apoptosis induced by UVB in the model, and it could be potentially inferred from corresponding biomarkers collected before the start of the treatment (*e*.*g*., number of apoptotic cells measured from biopsies 24-48 hours after localised delivery of phototherapy). Taken together with the above results, these data provide evidence that our model can simulate personalised response to UVB therapy, individual dosimetry and UVB administration regimes in clinical practice.

### 3.3 Representation of UVB response by the UVB sensitivity parameter enables fitting PASI trajectories

Using the patients’ UVB doses and their *full* PASI trajectories (including baseline PASI), we fitted the *uvb*_*s*_ parameter in our model to reproduce each patient’s PASI trajectory. These models are called *uvb*_*s*_-*personalised*, and in Figures 5a and 5b we show the PASI simulation computed by the *uvb*_*s*_- personalised models of two patients of our cohort. We fitted the *uvb*_*s*_ parameter for all our 94 patients. All of the derived *uvb*_*s*_ values were distributed between 0 and 1, median 0.24 and IQR range [0.16, 0.34]. The distribution (n=754) of the difference between the model simulated PASI Ψ^*^ (see Eq.(9)) and the patient’s actual PASI data is shown in Figure 5c. The resultant model simulations provided a close fit to the patients’ data with a mean PASI difference of 0.27 PASI units (recall that PASI ranges between 0 and 72).

**Figure 5:**
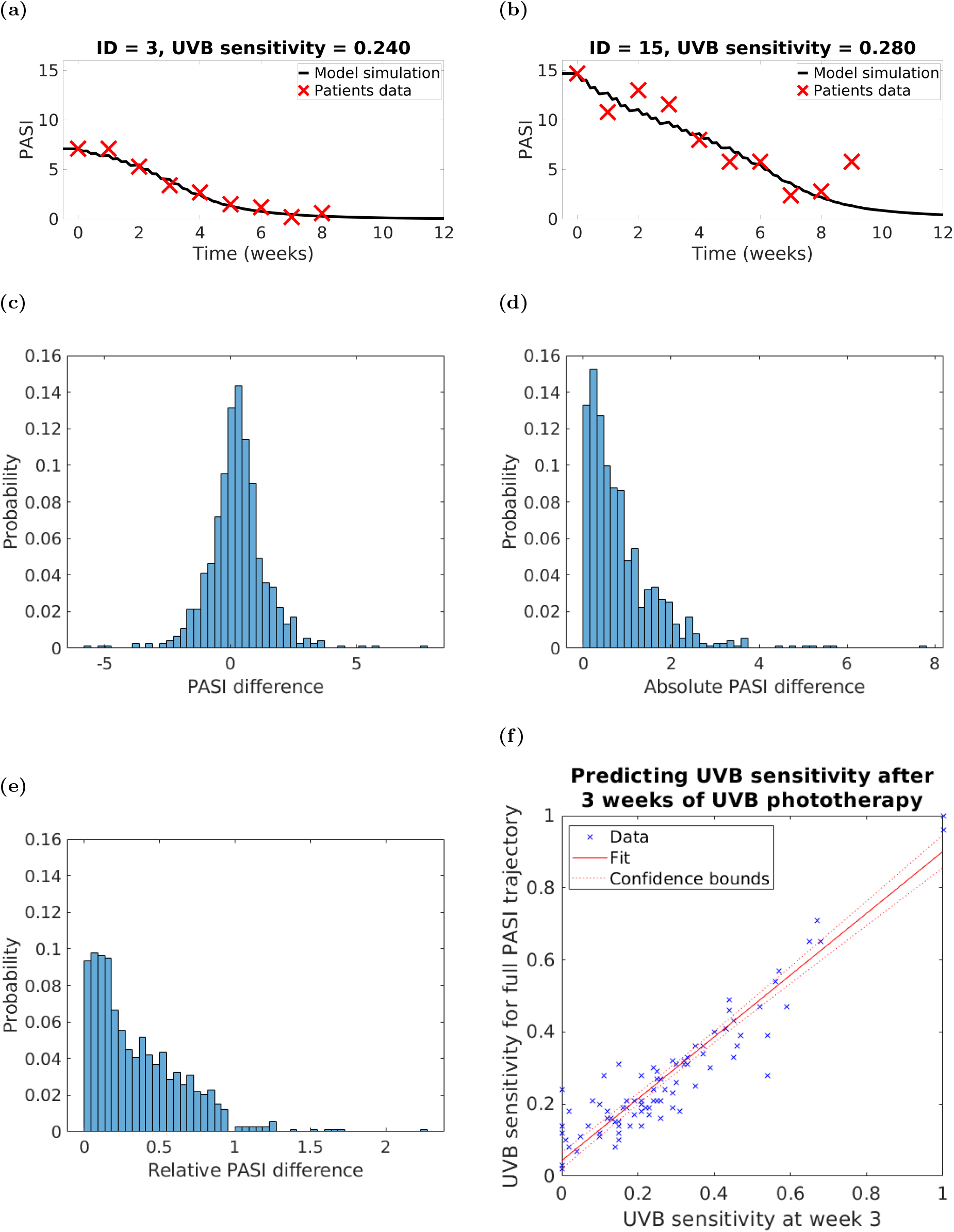
PASI measurements and UVB doses over the first three weeks of the therapy are sufficient to predict the UVB sensitivity *uvb*_*s*_ parameter, which allows high-accuracy model personalisation. Panels: (a,b) - results of parameter fitting for two different patients; (c) - distribution (n = 754) of the difference between the model PASI simulation Ψ^*^ and the patients’ actual PASI data; (d) - distribution (n = 754) of the absolute value of the difference between the model simulation and patients’ PASI data; (e) - distribution (n = 738) of relative PASI difference calculated as a ratio between the absolute PASI difference and the corresponding PASI value (excluding those values for which corresponding PASI = 0; n = 16), (f) - results of fitting *uvb*_*s*_ after 3 weeks with respect to fitting *uvb*_*s*_ at the end of the therapy.

Given a patient with a total of *t* weekly PASI readings, we calculated (for *i* = 1, …, *t*) the *absolute* (*i*.*e*., |*PASI* _*i*_ − Ψ^*^(7 · *i*)| = *AD* _*i*_, see Figure 5d) and the *relative*^1^ (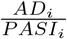, see Figure 5e) PASI differences and compared them to the PASI assessment discrepancies achieved by trained clinicians. For example, Ref. [22] reports on three formally trained physicians who performed 720 image-based PASI assessments of 120 patients with psoriasis (every physician assessed PASI twice: at week 0 and 4 weeks later). The mean and standard deviation (see Table 3) of the absolute and relative PASI differences of our models were very similar to the errors made by formally trained physicians assessing patients’ PASI.

**Table 3:** Simulation of PASI outcomes by our ODE model that are comparable to PASI assessments made by formally trained physicians and nurses. The *uvb*_*s*_ parameter of the models was estimated taking into account the full PASI trajectory.

| Source | Absolute PASI difference (mean±sd) | Relative PASI difference (mean±sd) |
| --- | --- | --- |
| Inter-rater difference [22] | 1.968 ± 2.480 | 24% ± NA |
| Intra-rater difference [22] | 1.984 ± 2.175 | 29.2% ± NA |
| Our model difference | 0.842 ± 0.838 | 35.4% ± 30.1% |

Together, these results suggest that given the patient’s baseline PASI, their UVB doses administered during the therapy and the UVB sensitivity parameter, our model can simulate PASI outcomes along the trajectory and final PASI that are indistinguishable from PASI assessments made by trained professionals. We note that the UVB sensitivity parameter could be estimated at baseline (*e*.*g*., by measuring the rate of apoptosis in skin biopsies) or early during the therapy, as we explain in Section 3.4. Therefore, our model can be used to predict individual patient response to phototherapy and enables design of personalised UVB regimes to improve outcomes, initially *in silico*, but ultimately in a clinical trial.

### 3.4 The UVB sensitivity parameter can be estimated at the third week of treatment

We found no statistically significant associations between the derived *uvb*_*s*_ values and the available patients’ clinical variables collected at baseline (*i*.*e*., BMI, age, sex, smoking status, alcohol consumption, skin type, baseline MED, age of psoriasis onset, previous phototherapies). Therefore, we tested whether the *uvb*_*s*_ parameter could be predicted by using only a portion of a patient’s PASI trajectory. We discovered that the earliest reasonable prediction (*R*^2^ = 0.895 and adjusted *R*^2^ = 0.894; root mean square error = 0.069, Figure 5f) was made by fitting *uvb*_*s*_ with the data available at the end of week 3 of the therapy, *i*.*e*., four PASI measurements and nine UVB doses. These data show that we can make a reasonable estimation of a patient’s UVB sensitivity parameter value by the end of the third week of the therapy which can then be used to predict subsequent response to UVB phototherapy. This discovery opens the path to personalisation of therapy.

### 3.5 The UVB sensitivity parameter and immune stimuli enable stratification of flaring patients

Flares are spontaneous worsening of a patient’s psoriasis symptoms and signs that can happen both on and off therapy. For example, Figure 5b illustrates an unexpected and sustained increase in a subject’s PASI (outside of observer error range [22]) despite ongoing UVB phototherapy. We hypothesised that our *uvb*_*s*_-personalised models could be used to distinguish between psoriasis flares and PASI assessment discrepancies. We remark that in case *uvb*_*s*_ is not available from clinical biomarkers, one could assume a baseline value or estimate a value after three weeks of therapy as previously discussed.

For each patient with *t* weekly PASI measurements we looked at their PASI errors, computed as

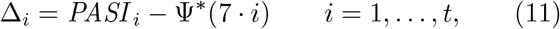

where Ψ^*^ is the *uvb*_*s*_-personalised model PASI simulation (see Section 2.3), and the relative PASI errors with respect to the model simulation, calculated as

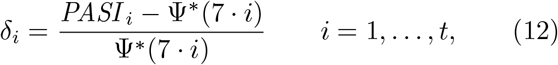

because unlike the patients’ PASI, the value of Ψ^*^(7 · *i*) is always strictly positive.

Utilising the DBSCAN clustering algorithm [20], we clustered the calculated PASI differences of all patients (n=754 PASI measurements) into two groups: PASI assessment errors and potential flares. We asked DBSCAN to identify two groups (see Figure 6a) within the data: the main cluster (black triangles) and the outliers (red crosses). By considering all positive relative error values (*i*.*e*., the *δ*’s) as flares (n=98) we fitted a *threshold curve*

**Figure 6:**
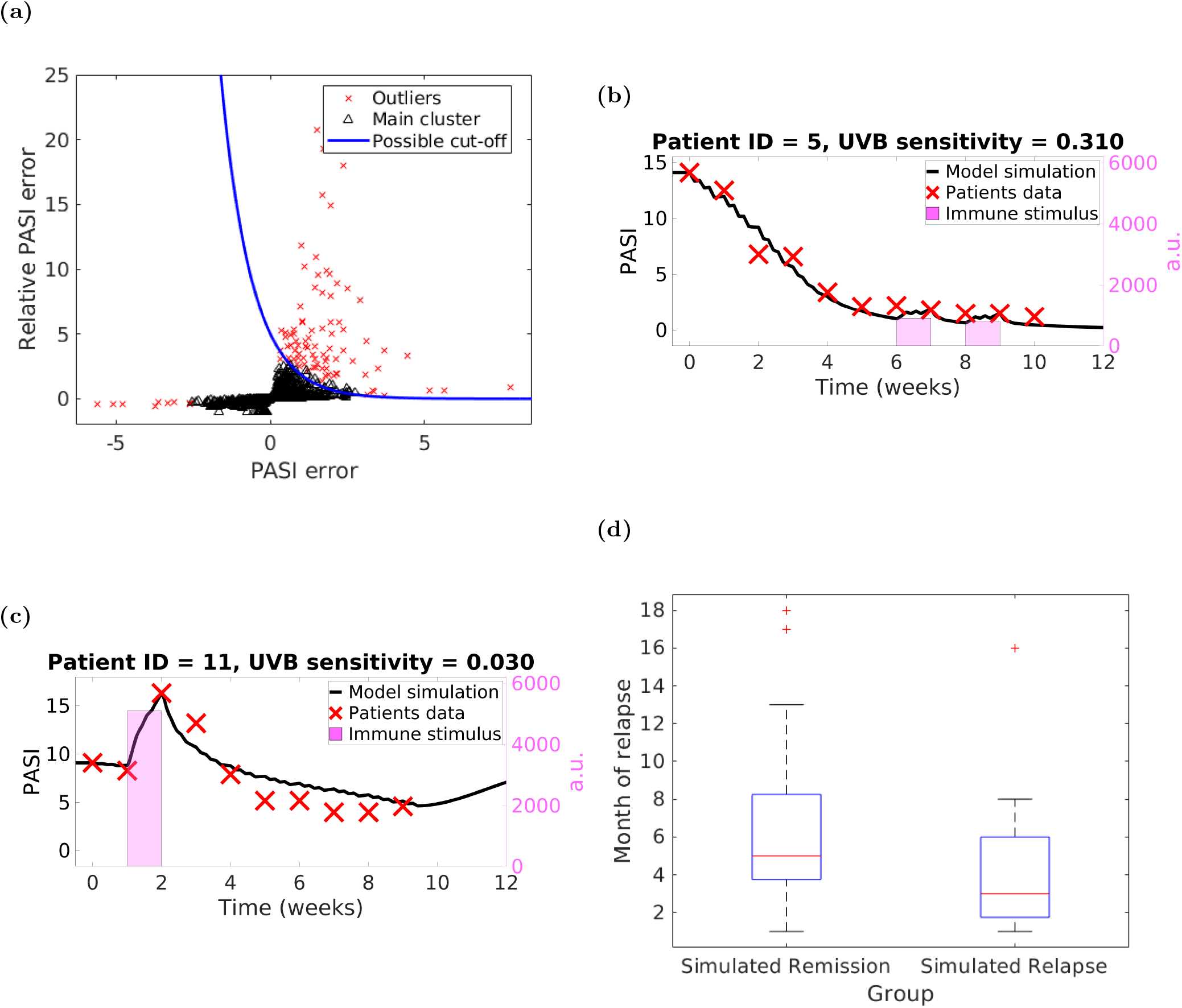
Our ODE model enables identification of psoriasis flares. Panels: (a) - distribution of relative PASI difference with respect to absolute PASI difference computed by DBSCAN with *ϵ* = 0.5 and minimum number of points 20 (red crosses – outliers, and black triangles – main cluster) and a possible curve separating flares (all the points above the blue line) and PASI measurement errors (all point below the line); (b,c) - examples of fitting immune stimuli for two different patients, (d) - patient groups based on model predictions of psoriasis relapse: our personalised flares-enabled model predicts Remission (n=49) and Relapse (n=13) groups.

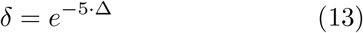

that separated the outliers (potential flares) from the main cluster (potential PASI assessment errors). (The 98 potential flares are distributed over 47 patients out of 94.)

We note that the threshold curve was identified using a large number of PASI measurements in which the sequential nature of the measurements was not retained. Therefore, this would enable using our *uvb*_*s*_- personalised models for ‘online’ detection of flares in the clinic as follows: every time a PASI measurement is obtained one computes both the simple (Δ) error and the relative (*δ*) error and then decides whether the pair sits above (flare) or below (measurement error) the threshold curve. Hence, as soon as a flare is detected the patient’s phototherapy regime can be modified (through increasing the frequency of irradiations and/or adding more UVB doses) to achieve a greater % improvement in PASI during therapy or a longer period remission.

The mean assessed PASI value was 8.8 and the range was [0.7, 34.8] (the corresponding figures in our dataset are 4.01 and [0, 31.8]), and the best mean absolute PASI difference was (reported for observers 1 and 2) 1.968 with standard deviation of 2.480, while the best mean absolute intra-observer PASI difference (the difference between the PASI assessments for observer 2) was 1.984 with a standard deviation of 2.175. NA – value is not available.

### 3.6 Flares during the therapy are associated with shorter remission

Our *uvb*_*s*_-personalised models provide evidence that detecting flares is extremely important not only for improving overall model fit, but also for predicting patients’ remission period.

To improve model fit, we applied an immune stimulus (iteratively increasing its strength starting from 0 while the sum of the squared errors (10) decreases) for 7 days for every week which is identified as a flare by the threshold curve (13). The resulting model is called *flare-enabled*. Figure 6b shows an example of a “flare” that was predicted and probably could not be detected visually. Our model implements a weak but persistent immunological stimulus but this result could also possibly be explained by increasing adaptation and resistance to UVB, which we do not currently model. In contrast, Figure 6c demonstrates a substantial flare in response to larger immunological stimulus (see Figure S5 for the model simulations without taking flares into account).

Next, we studied patients’ remission. We dropped therefore patients who were lost to follow-up (n=26) and those who did not relapse within the 18 month period (n=6; our model correctly predicts remission for all these patients). Then we divided the remaining patients (n=62) into two groups based on their simulated post-therapy PASI values: those whose *uvb*_*s*_-personalised, flare-enabled model simulations predicted relapse were assigned to a *Simulated Relapse* group (n=13), and the rest were placed into a *Simulated Remission* group (n=49). When compared to the actual, observed behaviour (see Figure 6d), the patients in the *Simulated Relapse* group demonstrated a shorter remission period (median value of 3 months *vs*. 5 months, Kruskal-Wallis p-value=0.035) compared to the *Simulated Remission* group. (*Simulated Relapse* group: mean = 4.23, standard deviation = 4.17, range = [1, 16] and IQR = [1.75, 6]; *Simulated Remission* group: mean = 6.45, standard deviation = 4.7, range = [1, 18] and IQR = [3.75, 8.25].)

We conclude that detecting flares is important so that therapy amendments can be applied not only to achieve better clearance at the end of phototherapy but also with the aim of extending the remission period.

## 4 Discussion

There is increasing interest in the application of systems biology modelling in medicine and immune-mediated inflammatory disorders [21]. To the best of our knowledge, the ODE-based model presented in this paper delineates for the first time the complex interactions between immune stimuli, keratinocyte growth, differentiation and apoptosis in the onset of psoriasis lesions, regulation of psoriasis flares and plaque resolution during UVB phototherapy. Our model features two stable steady states – non-lesional and psoriatic skin; switching between them occurs through immune stimuli and UVB phototherapy. We further include PASI modelling, which is necessary for a clinically valid model. Importantly, we demonstrate that individual patients’ PASI trajectories, recorded in response to UVB therapy, can be simulated by a designated model species and by estimating a single individualised model parameter (called “UVB sensitivity”) that is proportional to the rate of UVB-induced apoptosis. Notably, we show that the value of the UVB sensitivity parameter can be estimated by the end of the third week of a patient’s UVB therapy. We show that within a cohort of 94 patients this model enabled the prediction of individual patient outcome at the end of phototherapy based on baseline PASI, UVB dosimetry and the early trajectory of PASI response. The level of accuracy was within the error limits of PASI assessments made by trained professionals. Together these results support the prediction of longer term clinical outcomes that can be tested in the clinic.

Over the past twenty years several papers have reported agent-based models of human epidermis formation and homeostasis [52, 25, 2, 53, 41]. However, due in part to the disease complexity and to the difficulty of obtaining clinical data, there are few publications addressing the computational modelling of psoriasis and its treatment. An early work [26] proposed a 2D agent-based model of psoriasis formation that included keratinocytes only; and a psoriatic state was induced by manipulating the time transit-amplifying cells are allowed to proliferate for. Importantly this model did not include the concept of disease severity (*e*.*g*., by incorporating PASI), treatment response and was only qualitatively validated. An earlier work from our group introduced a 2D agent-based model that featured two stable steady states and modelled response to UVB phototherapy [59]. Whilst producing an interesting visualisation and quantitative readout of the psoriasis clearance process which provided insight into the mechanisms of clearance, the agent-based model was limited by a lack of personalisable data, including disease severity (PASI) and did not explicitly include immune cells, limiting its generalisability to other psoriasis treatments such as biologics. Recently, an ODE model [66] has been developed that proposes the interesting hypothesis that the epidermis phenotypes result from the homoeostasis of two distinct families of cells: healthy and psoriatic keratinocytes. While the model mostly behaves in a way consistent with the dynamics of psoriasis, the hypothesis on which it rests has not found confirmation in the medical field, to the best of our knowledge [49]. Furthermore, the UVB phototherapy regimes used for clearing psoriasis seem unrealistically short (seven doses over 16 days *vs*. 25-30 doses, 3x weekly in common clinical practice), which would likely entail high erythemogenic doses.

With respect to the use of machine learning approaches for psoriasis treatment, a recent paper [24] has employed unsupervised machine learning techniques to identify subgroups in patients undergoing biologic treatments based on their PASI trajectory over time. The analysis has revealed a model with four classes of response trajectories with distinct clinical outcome and remission. However, it is unclear whether the model and its class characterisation are powerful enough to predict individual patient outcome. In a recent paper [57], we report the development of a machine learning approach combined with a logistic regression model to predict final PASI using the first 2-3 weeks of PASI measurements during UVB phototherapy. Thus, compared to previous publications, notable advances of our model reported herein are the flexibility to be personalisable at the individual subject level with a single parameter (UVB sensitivity), the possibility to include flares during the therapy (via an extra parameter mimicking a transient immune stimulus), and the ability to capture the PASI dynamics during phototherapy in a way indistinguishable from actual PASI measurements in clinical practice. These are crucial factors that facilitate the clinical application of our model and represent a significant advance compared to the works surveyed above. Furthermore, the inclusion of key components of the immune system in our model enable its generalisation to biologic therapies, which are used for treating more severe forms of psoriasis. In particular, developing machine learning models predictive of psoriasis outcome for biologics is likely to be challenging due to the time-sparsity of data during the early phases of biologics treatments, in part related to the time frame of clinical follow up. As such, mechanistic models like ours built from both clinical data and experimental data within the literature will likely be instrumental for modelling biologics treatments.

Our model is based on three main assumptions: 1) apoptosis induced by a single UVB dose lasts for 24 hours and affects proliferating keratinocytes and immune cells equally (see Section 2.2; based on our previous studies [59]); 2) cell growth depends on cell density (this is a common assumption in modelling – more details are given in Section SM.5); and 3) our PASI model does not take the disease area into account, since our model tracks cell densities only, and in most cases the PASI subscores are seldom recorded in the clinic. Additionally, our model focuses on whole body phototherapy of mild to moderate psoriasis and whether more severe psoriasis responds clinically in a similar manner remains to be determined experimentally.

With respect to strengths, our model is computationally efficient: simulations generally take only a few seconds on a standard desktop or laptop computer. Furthermore, with only two tuneable parameters – UVB sensitivity and immune stimulus – our model is able to fit with high accuracy real PASI data (in the sense that PASI outcome model simulations are comparable to PASI measurements by clinicians), including flares during the therapy. The UVB sensitivity parameter could be estimated at baseline by clinical biomarkers or at week 3 during UVB phototherapy by simple PASI readings (*R*^2^ = 0.895 and adjusted *R*^2^ = 0.894; RMSE = 0.069). Our model supports personalised therapy outcome prediction by being able to include UVB doses administered in the clinic and different delivery patterns (*i*.*e*., 3x weekly *vs*. 5x weekly). Finally, our model is highly generaliseable: it already has the foundations necessary to accommodate other therapies (*e*.*g*., biologics) that block the immune stimulus (*e*.*g*., anti-IL-17 or anti-IL-23 antibodies) or induce growth arrest.

As for limitations, it is worth noting that the PASI errors clustering can vary depending on the utilised clustering algorithm and its hyper-parameters. This underscores the need for further experimental work to identify the biomarkers that are associated with disease flares and such studies can now be guided by our computational model. Although not strictly a limitation of our modelling, one should keep in mind that PASI does not provide an objective, completely reliable assessment of psoriasis severity. By definition, it is at least in part observer dependent, and this could lead to significant differences between predicted and observed outcomes in the clinic. Finally, our model provides a broader picture of the compartments within epidermis, and therefore, it does not take into account certain specific spatial details. For example, it cannot distinguish between the actively dividing and the dormant stem cells and transit-amplifying cells. Also, cell differentiation is modelled as a single step process while in reality it involves multiple stages.

The current ODE model is also confined by its transition between two main steady states. Thus, the initial psoriasis state prior to UVB phototherapy is independent of individual PASI scores, and therefore the cell densities at time 0 are equal. In expanding our model to consider moderate to severe psoriasis and its response to biologics, it would prove highly complex and computationally challenging to model individuals according to their exact baseline PASI. We will thus explore broad categories of baseline psoriasis activity and how this influences response (for example, PASI 5 to 10, PASI greater than 10, PASI greater than 20).

Our model is based on ODEs and therefore it necessarily needs to abstract some cellular mechanisms to give an ‘overall’ picture of the system dynamics. For example, modelling the different modes of division of stem cells is not currently readily compatible with ODEs. Ideally, the state of every cell type should be represented by a different species (*e*.*g*., TA cells that underwent one and two cycles of division will be modelled by two different species. Similarly, symmetric and asymmetric divisions should be handled as separate ODEs). This would significantly complicate the model and our approach of relating cell species and the clinical outcome of the therapy in the form of PASI, which is crucial for personalised treatments. Agent-based models would be a better framework to study in detail this kind of cellular mechanisms, although at a much heavier price in terms of model construction and computational effort. (The latter in particular would prevent such a model from being used in ‘real time’ in the clinic, while ODEs simulation is nearly instantaneous.) Building on our previous work [59], we are further developing a 3- dimensional agent-based model of psoriatic skin and UVB therapy that will be well suited to exploring symmetric and asymmetric divisions and other important issues arising from experimental studies [49]. Although biologic therapy has not been considered in the current work, the explicit representation of immune cells (DC, T) and mediators (TNF, IL-17, IL-23) opens up the possibility of simulating the effects of biologics by adapting the current model. In addition, we suggest that further studies to identify biomarkers (*e*.*g*., obtained from blood or RNA-sequencing of biopsies) associated with the proposed UVB sensitivity parameter should be conducted. Data from further clinical studies should also be used to refine and improve the model parameters. Finally, validating the model in a clinical setting will open up its use for personalised treatment of psoriasis in practice.

In conclusion, our computational model of psoriasis explicitly describes the interaction between the immune system and epidermal keratinocytes in transitioning between the steady states of lesional and non-lesional psoriatic skin. Importantly our model underscores the importance of apoptosis as an important mechanism in clearance of psoriasis in response to UVB. Together with data from our experimental studies [59, 1] this suggests that targeting of apoptosis in drug development and therapeutic compound screening may prove useful. Additionally, our model distinguishes disease flares, and supports prediction of amending individual UVB phototherapy regimes based on the patient’s initial response that include for example personalised delivery schedules (*i*.*e*., 3x weekly *vs*. 5x weekly phototherapy). Therefore, our work represents a crucial step towards precision medicine for psoriasis.

## Code and Data Availability

Matlab code for our model is available at https://github.com/pzuliani/psoriasis/.

## Author Contributions

F.S. developed the model, ran the simulations, analysed the results and wrote a draft manuscript; G.R.S. contributed improvements to the model; N.J.R. and S.C.W. provided the clinical data; P.Z. and N.J.R. conceived the project, supervised the research and contributed to writing the manuscript. All authors discussed the results and approved the final manuscript.

## Acknowledgments

F.S. was supported by a Rosetrees Trust award (M651) while at Newcastle University. This study received funding from the National Institute for Health Research (NIHR) Newcastle Biomedical Research Centre, the British Skin Foundation, the British Phototherapy Group. N.J.R. is also supported by the Newcastle NIHR Medtech and In Vitro Diagnostics Co-operative and is a NIHR Senior Investigator. We are grateful to the research and phototherapy nurses who helped throughout this study and to the patients who participated.

## SM Supplementary Material

### SM.1 Model details

We used the SMT solver dReal to show that there are only 3 balls of diameter at most 3 · 10^−9^ in the domain [10^−9^, 10^32^] × … × [10^−9^, 10^32^] that can contain the solutions. (Although this does not formally guarantee that there are at least 3 solutions.) So if there are more than one solution within each such ball (there are 32 solutions overall), they are at most 3 · 10^−9^ far from each other. Below are the intervals where the solutions for each species might be. So basically there are no solutions in the regions ±10^−9^ from the right and the left bounds of the intervals below.

Solution intervals group 1 (healthy state):

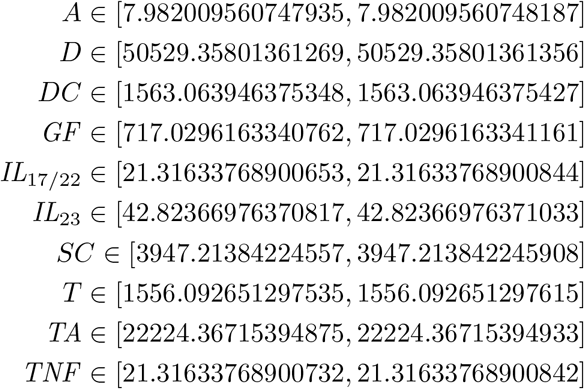

Solution intervals group 2 (transition state):

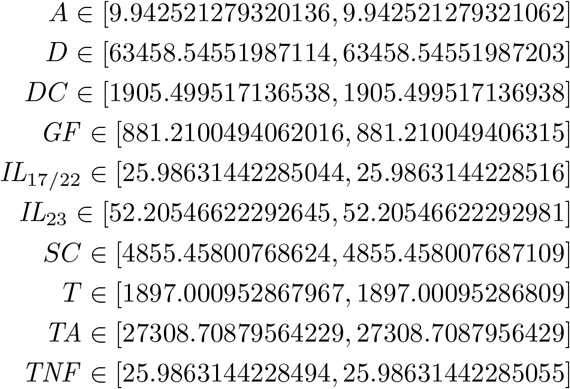

Solution intervals group 3 (psoriatic state):

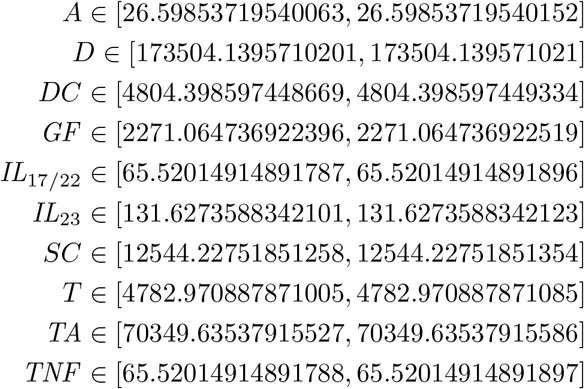

### SM.2 Bimodal behaviour of the model

Our model presented in Figure 1b and equations (1) features bimodal behaviour with two stable positive real (*i*.*e*., “healthy” and “psoriatic”) steady states and one unstable positive real (*i*.*e*., “transition”) steady state (see Table 2 for the corresponding cell densities in the three states). The transition from “healthy” to “psoriatic” state happens via an introduction of a sufficiently strong immune stimulus (this causes an immediate increase in the number of dendritic cells), while the transition from “psoriatic” to “healthy” state can be achieved through inducing a sufficient amount of apoptosis in the proliferating keratinocytes and the immune cells. In Figure S1 we show phase planes demonstrating the behaviour of the model depending on the initial density of dendritic cells and transit-amplifying cells, which are the two key species involved in the transition between the steady states.

### SM.3 Parameter sensitivity analysis

To evaluate the robustness of our model with respect to rounding changes in key parameter values, we performed a sensitivity analysis for the model parameters from Table 1 whose values feature more than two significant digits (*i*.*e*., *sc*_2_, *sc*_2*ta*_, *sc*_20_, *ta*_2_, *ta*_20_, *ta*_2*d*_, *d*_20_ and *d*_*desq*_). We analysed steady states for 256 combinations of these parameters by rounding their values up and down to two significant digits. (Specifically, we used MATLAB to find all the zeroes of the right-hand side of the ODE system (1), initialised with each of the 256 combinations.) As a result, our model features bistability (*i*.*e*., presence of two stable steady states; see Figure S2) for 200 parameter combinations, while in the remaining 56 cases bistability was broken (*i*.*e*., the model featured only one steady state). The results show that our model is generally robust with respect to small variations of the parameter studied. We note that the higher value of *d*_*desq*_ is always necessary to break bistability. In addition, the yellow column suggests that *d*_*desq*_ and *ta*_2*d*_ are the two major ‘culprits’.

**Figure S1:**
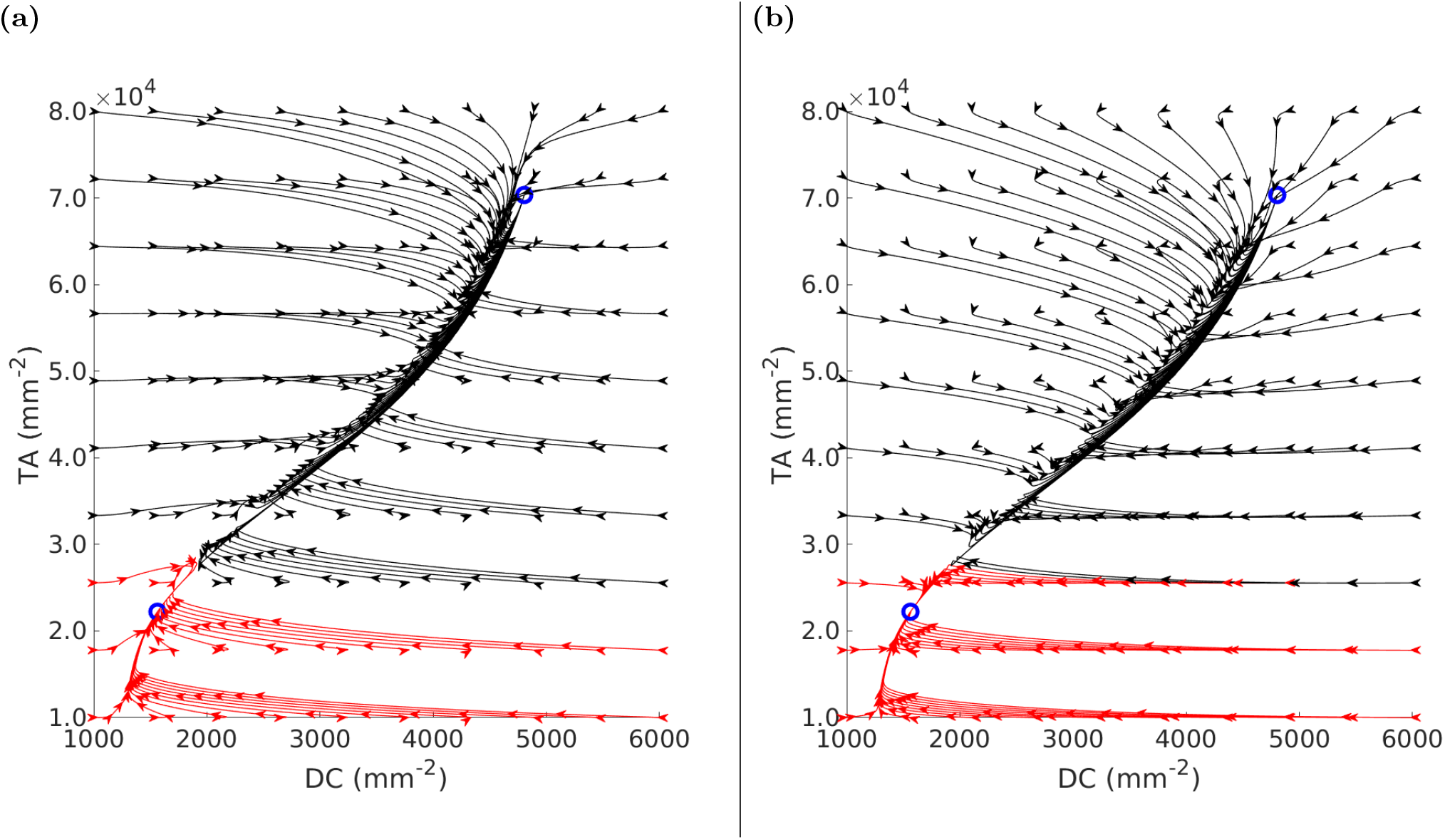
Phase planes demonstrating the effect of changing initial values of *DC* and *TA* species on converging to the two stable steady states of the model. Red and black arrows represent model trajectories converging to the healthy (lower blue circle) and the psoriatic steady state (upper blue circle), respectively. Panels (a): all the remaining species were initialised to the healthy steady state; (b) remaining species initialised to the psoriatic steady state.

### SM.4 Growth and differentiation rate of keratinocytes

The growth rate and the differentiation r ate o f keratinocytes are the positive terms of the differential equations (see (1)) for cell species *SC, TA*, and *D*. The terms are (*TNFα* + *IL*_17*/*22_)(*sc*_2_*SC* + *sc*_2*ta*_*SC* + *ta*_2_*TA*) and *ta*_2*d*_(*TNFα* + *IL*_17*/*22_)*TA* for the growth rate and differentiation r ate, r espectively. Thus, the change of keratinocytes growth rate in the two stable states of the model (healthy and psoriasis) can be calculated as the ratio:

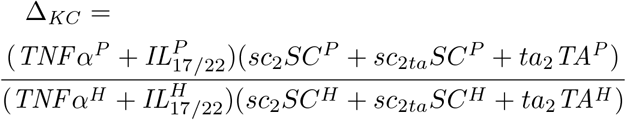

**Figure S2:**
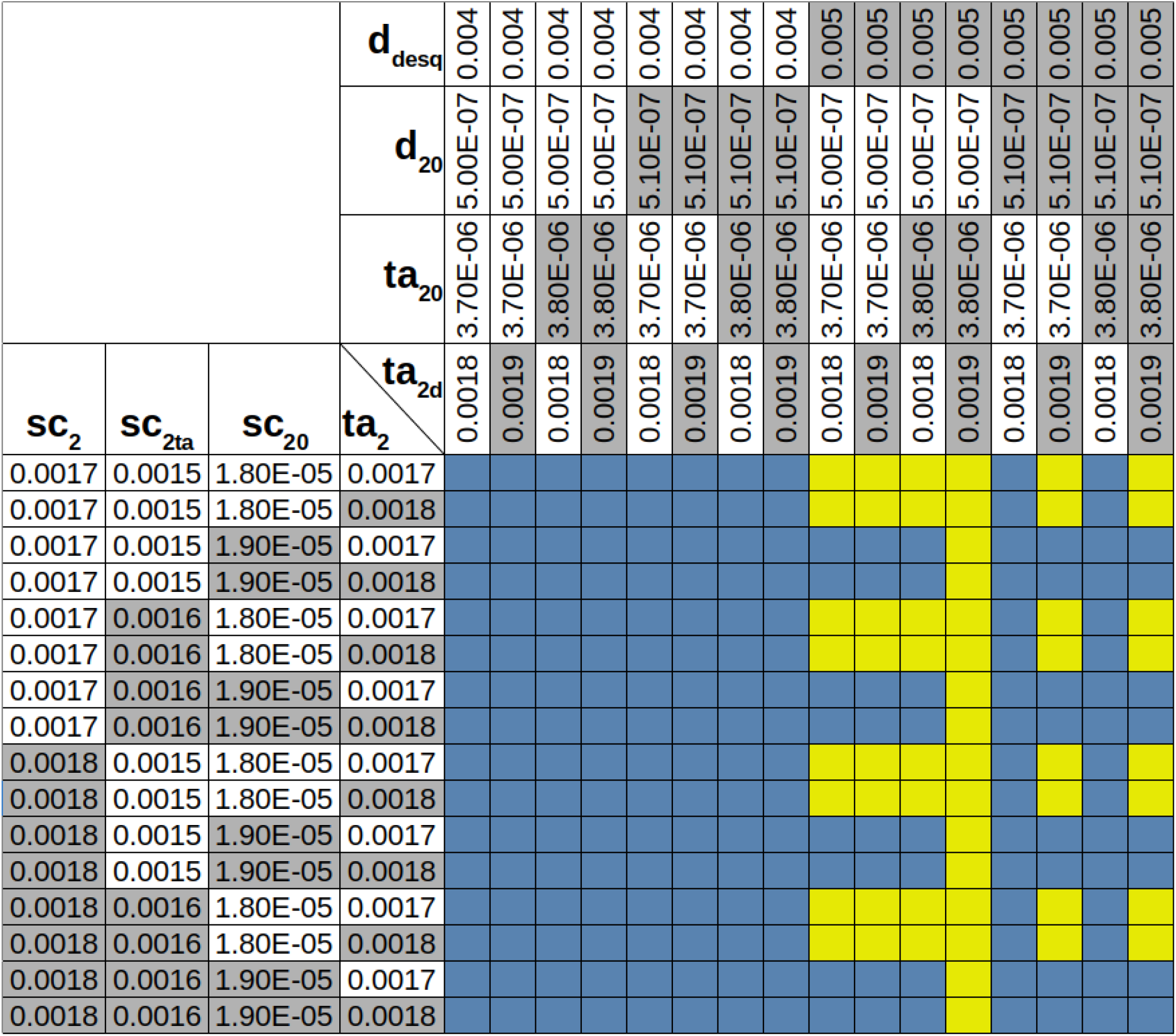
Parameter sensitivity analysis. The table shows the effect of changing the model’s parameters’ values on the number of steady states of the model (cells with grey and white backgrounds contain the upper and the lower parameter values, respectively). Blue cells denote parameter combinations for which bistability holds, while yellow cells denote parameter combinations for which bistability breaks (*i*.*e*., the model features only one real positive steady state).

where the superscripts *P* and *H* represent the values of the model species in the psoriatic and the healthy steady states, respectively. By utilising the values found in Table 2 we compute Δ_*KC*_ ≈ 9.74. Similarly, the change in the differentiation rate is

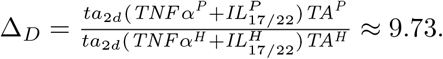

### SM.5 Keratinocytes growth *vs*. epidermis cell density

In our model we assume that cell growth rates depend on the cell density. This assumption was made from observing the dependency between epidermal turnover time and epidermal thickness. Turnover time is the amount of time needed to replace all cells in a particular compartment. It can be calculated as the total number of cells in the compartment divided by the rate of cell growth (cell death for systems in homeostasis). It has been reported that the thickness of psoriatic epidermis increases by around 2-5 times [59, SM], while the epidermal turnover time decreases by up to 4 times [59, SM]. (The interested reader may find more references in Section 1.5, 6.3.1 and 6.3.2 of [58].)

Suppose 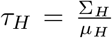 and 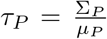 (where Σ_*H*_ and Σ_*P*_ are total cell densities in healthy and psoriatic epidermis, and *μ*_*H*_ and *μ*_*P*_ are the rates of cell growth in healthy and psoriatic epidermis) are the turnover times for healthy and psoriatic epidermis, respectively. Typically, the growth rate is described by the law of mass action as *k* · Σ, where *k* is the growth rate constant. Essentially, it states that the cell growth rate depends linearly on the cell density. However, we will show why the law of mass action is not directly applicable in our model without making further assumptions (*e*.*g*., growth limiting factors, forces, viscosity, limited nutrition, etc.).

Let 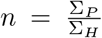 be the ratio between the thickness of psoriatic and healthy epidermis, and let 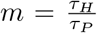 be the turnover time ratio between psoriatic and healthy epidermis. From the above it follows that *μ*_*P*_ = *m* · *n* · *μ*_*H*_, which means that epidermal thickness increases *n*-fold while the cell growth rate increases by *m* · *n*. This suggests that the growth rates *μ*_*P*_ and *μ*_*H*_ cannot be modelled by the law of mass action as we derive the following contradiction Σ_*P*_ = *n* · Σ_*H*_ (from the thickness ratio formula) and Σ_*P*_ = *m* · *n* · Σ_*H*_ (from the growth rates equation).

However, if we assume that the cell growth rate is defined by a nonlinear function *μ* = *k* · Σ^*p*^, where *p* > 1, we get the following system of equations Σ_*P*_ = *n*· Σ_*H*_ and 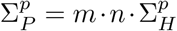. Solving it for *p* yields *p* = (log_*n*_ *m*) + 1. Thus, we model keratinocyte growth using the following ODE

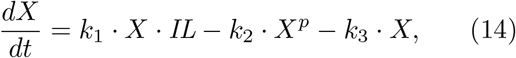

where the term *k*_1_ · *X* · *IL* describes the growth rate of keratinocytes as a law of mass action with *X* being the keratinocyte species and *IL* being the cytokine species, *k*_3_ · *X* represents cell degradation also modelled by the law of mass action, and *k*_2_ · *X*^*p*^ describes the limiting factors of keratinocyte growth due to an increase in cell density (this could be caused by nutrients limitation and spatial constraints, for example); we call *p* the “growth limiting constant”. (We note that this approach, also known as *logistic growth* because of the RHS of Eq. (14), has been in use since the 19*th* century for modelling growth dynamics that are subject to constraints, such as for example in ecological modelling with the well-know Lotka-Volterra equations for a predator-prey system.)

We emphasise that Eq. (14) is also used for differentiating keratinocytes (*D*): Eq. (1) for *D* features a *D*^*p*^ term. While in the equation for stem and TA cells the corresponding term is needed for bistability, the rationale for *D* is to enforce a decrease in transit time (turnover time for *D* species) while maintaining the same cell densities proportions in the psoriatic state. In addition, we assume that *n* = *m* (and hence, *p* = 2) in our model for simplicity. However, the epidermal thickness and the cell growth rate can be changed using parameters *n* and *m*, which provides modelling flexibility.

### SM.6 Apoptosis and Desquamation Rates

The dynamics of apoptotic cells (*A*) is governed by the equation 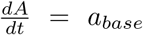(*SC* + *TA* + *DC* + *T* + *D*) − *a*_20_*A*. It is assumed that in healthy epidermis all cells undergo apoptosis at the same rate (relative to their cell mass). The rate of degradation of apoptotic cells is governed by the term *a*_20_ *A*, hence their mean lifetime is 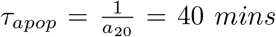 ([58, Figure 5-5] reports a median time of 20 minutes from an *in vitro* study, [29, Figure 1] states 1-3 hours for lymphocytes and up to 48-72 hours for keratinocytes), and [38, Figure 1d] demonstrates an apoptotic cell being destroyed within 20-40 minutes in mice). The apoptosis rate in terms of number of apoptotic cells per 1,000 can be calculated·as 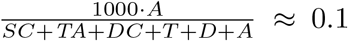 (the median number of apoptotic cells per 1,000 is reported to be 0.1 in untreated psoriatic skin [59, Figure 1b]). The epidermal desquamation rate in the model can be calculated as *d*_*desq*_ · *D*, and in the healthy state it is *d*_*desq*_ · *D*_*H*_ ≈ 240.16 *cells/*(*day* · *mm*^2^), where *D*_*H*_ is the density of differentiated cells in the healthy steady state. A similar figure for human skin is reported by [39].

### SM.7 Supplementary figures

Figures S3a, S3b and S3c demonstrate three different simulation scenarios where the same 30 UVB doses are administered once per week, but different UVB sensitivity values are simulated. It can be seen that UVB sensitivities of 0.05 and 0.1 do not induce psoriasis clearance, while a UVB sensitivity of 0.15 (*i*.*e*., three times the base value) does clear psoriasis. However, we remark that this is a purely hypothetical scenario as patients with different UVB sensitivities are very unlikely to be assigned the same UVB doses. Most likely a patient with higher UVB sensitivity will receive lower UVB doses.

**Figure S3:**
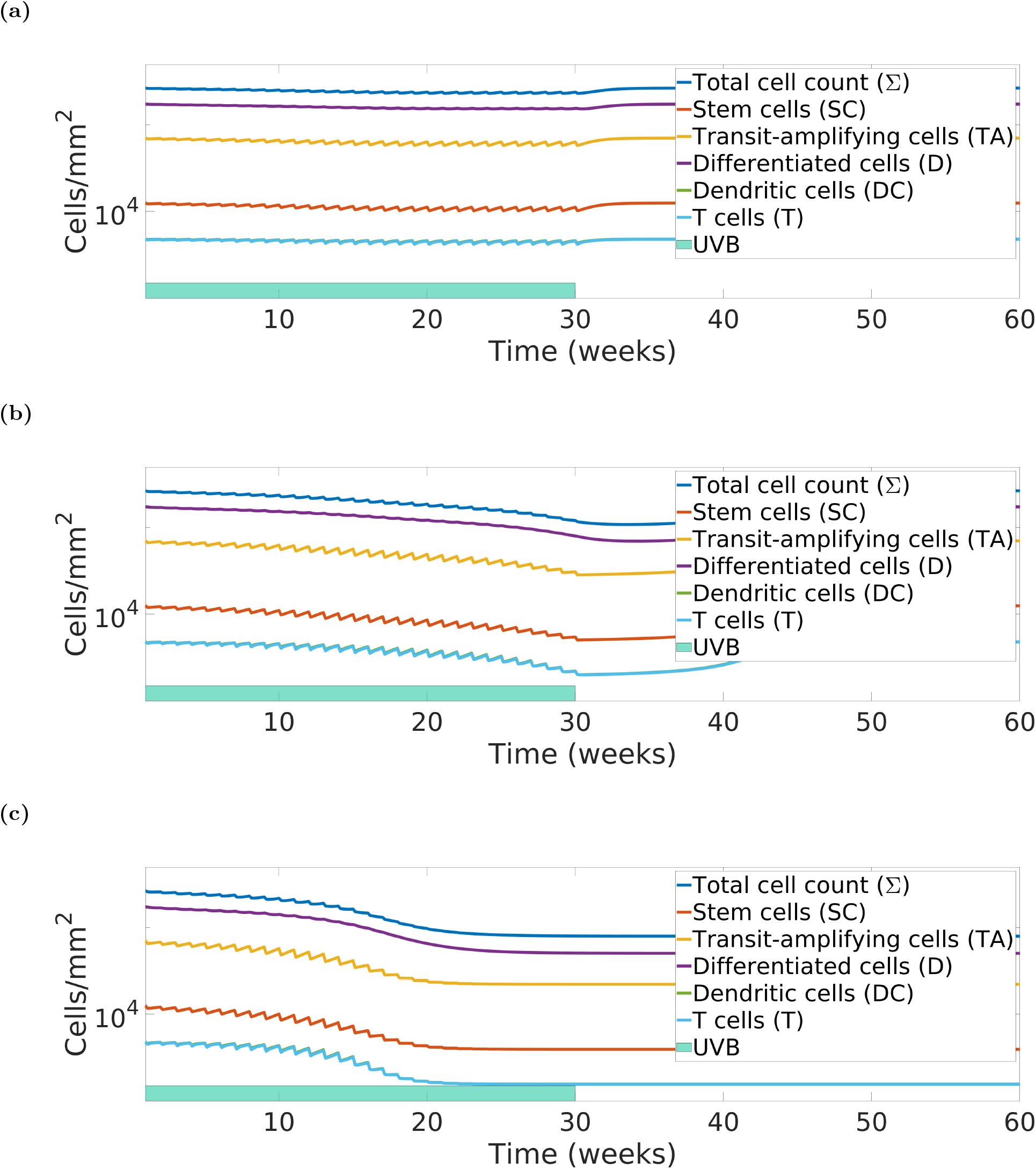
Model simulations of 1x weekly UVB phototherapy (30 doses) *vs*. UVB sensitivity. Panels: (a) *uvb*_*s*_ = 0.05: no clearance; (b) *uvb*_*s*_ = 0.1: no clearance; and (c) *uvb*_*s*_ = 0.15: psoriasis cleared.

**Figure S4:**
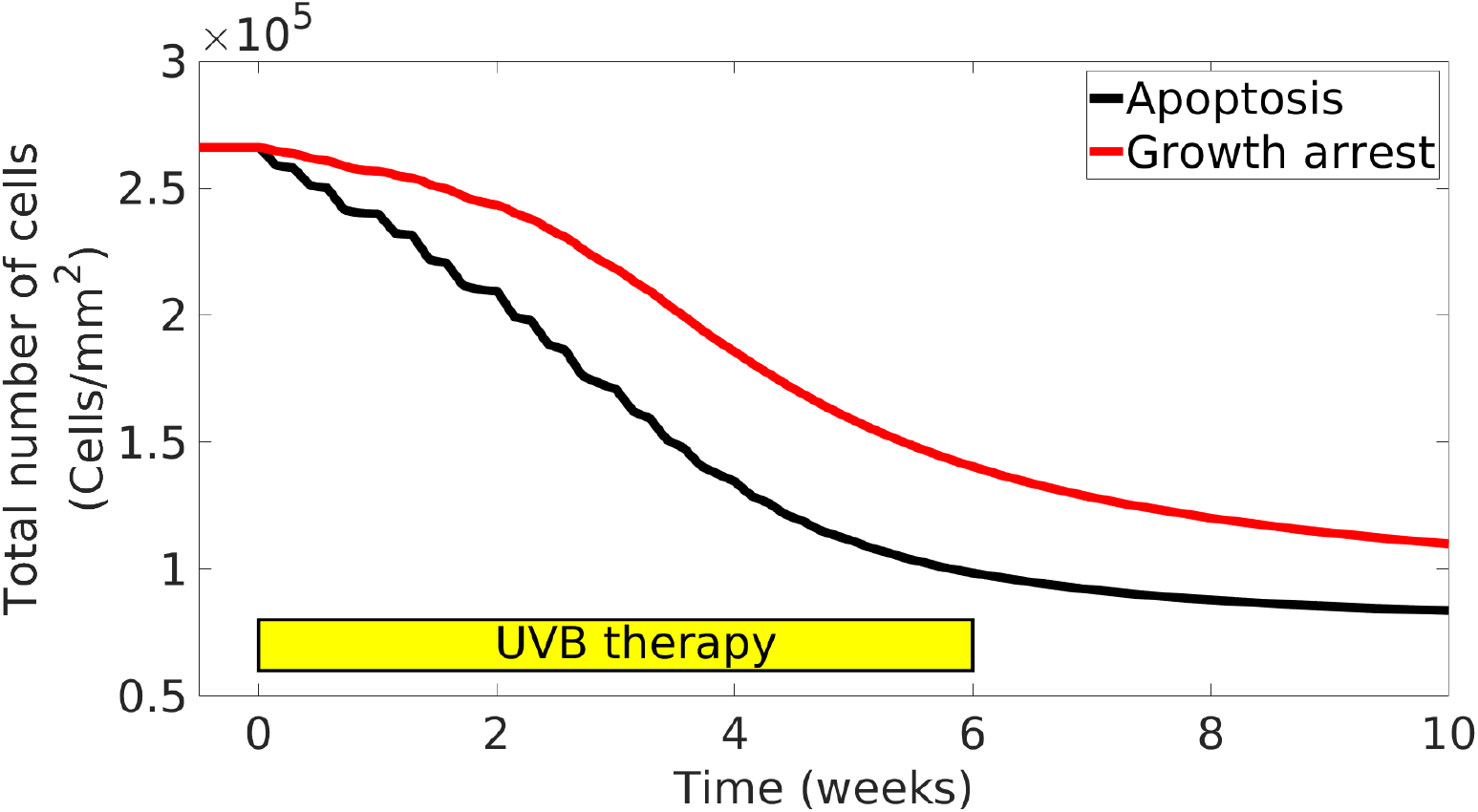
Model simulation of UVB-induced cell apoptosis *vs*. cell growth arrest. The Apoptosis model reaches the healthy steady state at the end of the UVB therapy, while the Growth arrest model is still in moderate psoriasis at the end of the therapy and an eventual psoriasis clearance. The latter contradicts clinical observations, and it was thus disregarded.

**Figure S5:**
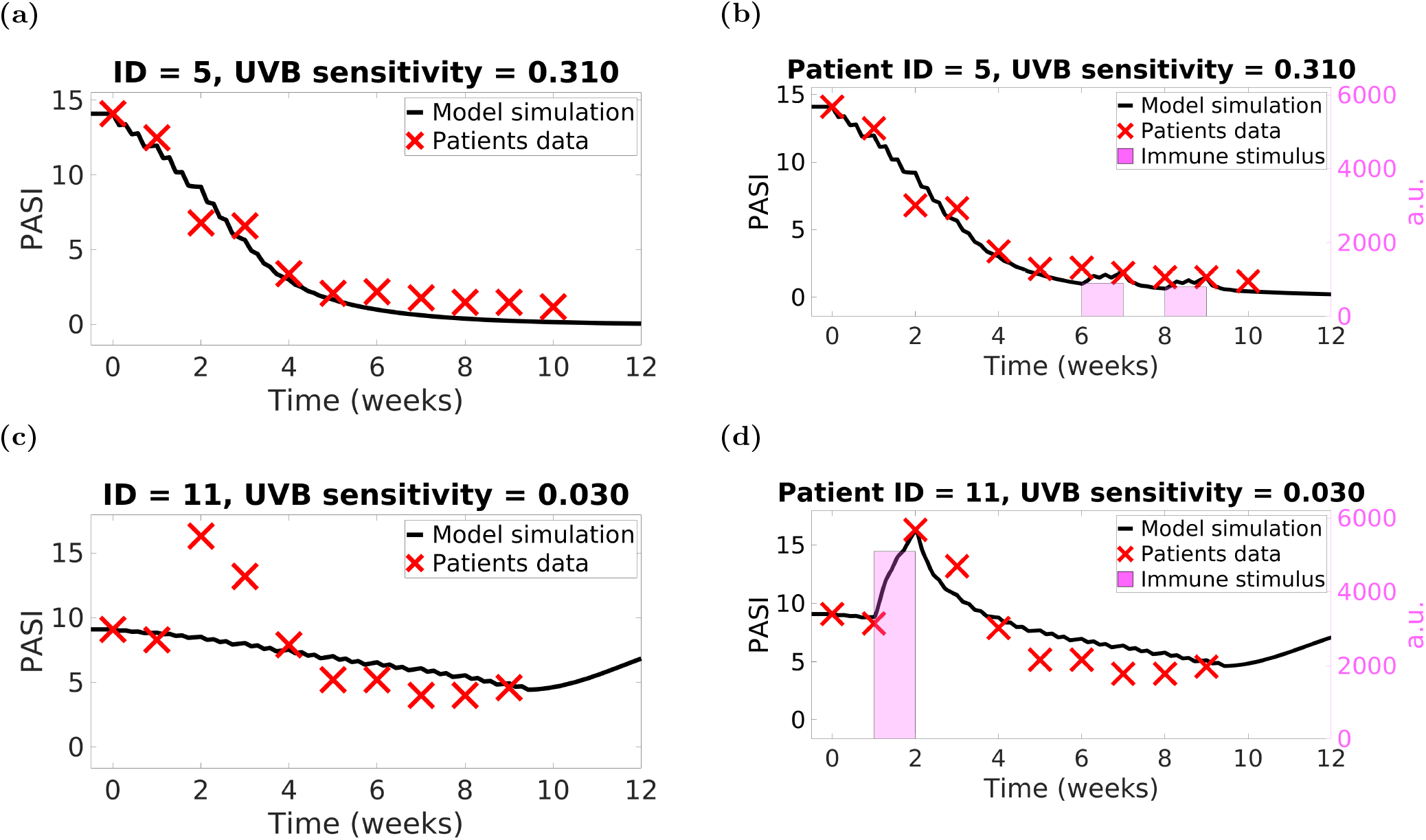
Modelling flares by introducing an immune stimulus to the model. Panels (a), (c): two patients PASI trajectories and their model simulation without immune stimulus. Panels (b), (d): the same two patients with their respective model fitted with immune stimuli, allowing a much closer match between patient data and model simulation. Panels (b) and (d) are the same as Figure 6b and Figure 6c, respectively: they are reported here for the reader’s convenience.

## Footnotes

1 Here we only used the values whose corresponding PASIs were non-zero (n = 738). In the remaining (n=16) cases we obtained the following distribution of absolute difference: median = 0.33, mean = 0.42, standard deviation = 0.37, range = [0.11, 1.67] and IQR = [0.24, 0.44].

